# Novel CRISPR-based detection of *Leishmania* species

**DOI:** 10.1101/2022.04.29.490093

**Authors:** Eva Dueñas, Jose A. Nakamoto, Luis Cabrera-Sosa, Percy Huaihua, María Cruz, Jorge Arévalo, Pohl Milón, Vanessa Adaui

**Author notes:** Corresponding author: Vanessa Adaui. Laboratory of Protein Evolution, Department of Experimental Medical Science, Lund University, Lund, Sweden (Jose A. Nakamoto). Laboratorio de Malaria: Vectores y Parásitos, Laboratorios de Investigación y Desarrollo, Facultad de Ciencias y Filosofía, Universidad Peruana Cayetano Heredia, Lima, Peru (Luis Cabrera-Sosa).

## Abstract

Tegumentary leishmaniasis, a disease caused by protozoan parasites of the genus *Leishmania*, is a major public health problem in many regions of Latin America. Its diagnosis is difficult given other conditions resembling leishmaniasis lesions and co-occurring in the same endemic areas. A combination of parasitological and molecular methods lead to accurate diagnosis, with the latter being traditionally performed in centralized reference and research laboratories as they require specialized infrastructure and operators. CRISPR-Cas systems have recently driven innovative tools for nucleic acid detection that combine high specificity, sensitivity and speed and are readily adaptable for point-of-care testing. Here, we harnessed the CRISPR-Cas12a system for molecular detection of *Leishmania* spp., emphasizing medically relevant parasite species circulating in Peru and other endemic areas in Latin America, with *L*. (*Viannia*) *braziliensis* being the main etiologic agent of cutaneous and mucosal leishmaniasis. We developed two assays targeting multi-copy targets commonly used in the molecular diagnosis of leishmaniasis: the 18S ribosomal RNA gene (18S rDNA), highly conserved across *Leishmania* species, and a region of kinetoplast DNA (kDNA) minicircles conserved in the *L*. (*Viannia*) subgenus. Our CRISPR-based assays were capable of detecting down to 5 × 10^−2^ (kDNA) or 5 × 10^0^ (18S rDNA) parasite genome equivalents/reaction with PCR preamplification. The 18S PCR/CRISPR assay achieved pan-*Leishmania* detection, whereas the kDNA PCR/CRISPR assay was specific for *L*. (*Viannia*) detection. No cross-reaction was observed with *Trypanosoma cruzi* strain Y or human DNA. We evaluated the performance of the assays using 49 clinical samples compared to a kDNA real-time PCR assay as the reference test. The kDNA PCR/CRISPR assay performed equally well as the reference test, with positive and negative percent agreement of 100%. The 18S PCR/CRISPR assay had high positive and negative percent agreement of 82.1% and 100%, respectively. The findings support the potential applicability of the newly developed CRISPR-based molecular tools for first-line diagnosis of *Leishmania* infections at the genus and *L*. (*Viannia*) subgenus levels.

## INTRODUCTION

Leishmaniasis is a vector-borne disease of global public health importance caused by intracellular protozoan parasites of the genus *Leishmania*. The disease affects 12 million people spread in 98 countries, with approximately 1.6 million new cases occurring annually (Alvar et al., 2012). In leishmaniasis endemic areas, asymptomatic infections by *Leishmania* are common (representing ∼10%-60%; Bañuls et al., 2011; Ibarra-Meneses et al., 2022), yet some infected individuals develop a variety of clinical manifestations that affect the skin and/or mucosal tissues (tegumentary leishmaniasis, TL) or internal organs (visceral leishmaniasis, VL) (Burza et al., 2018). In the Americas, TL is widespread, with 54,000 cases reported annually in 18 endemic countries (PAHO/WHO, 2020; Ruiz-Postigo et al., 2021). It encompasses skin lesions (96% of reported cases) and mucosal lesions (4% of cases) that can lead to permanent scars and devastating life-threatening mutilation of the nasopharynx, respectively (PAHO/WHO, 2019). These disease phenotypes are caused by different *Leishmania* species and are associated with diverse host-parasite interactions and human host immune responses (Carvalho et al., 2012; Gollob et al., 2014). Among the diverse Neotropical *Leishmania* species of the *Viannia* and *Leishmania* subgenera that circulate in endemic areas, the former are the most frequent cause of TL (Davies et al., 2000; Reithinger et al., 2007; Brito et al., 2009). Particularly, infections with *L*. (*Viannia*) *braziliensis* are predominant and of public health concern because of the associated risk of disease progression from cutaneous to mucosal leishmaniasis (Llanos-Cuentas et al., 1984; Marsden et al., 1986) and treatment failure (Arévalo et al., 2007; Adaui et al., 2016).

Early and accurate diagnosis of leishmaniasis is critical for its clinical management and timely treatment. Differential diagnosis of TL is challenging due to the clinical pleiomorphism and thus combines clinical characteristics, epidemiological factors, and multiple laboratory diagnostic tests because no perfect reference standard test exists (Adams et al., 2018). Screening for *Leishmania* amastigotes by microscopic examination of Giemsa-stained slide smears of human skin lesions and isolation of parasites from lesions in *in vitro* culture are mainly used for routine diagnosis of TL. However, these methods have suboptimal sensitivity (Weigle et al., 1987; Weigle et al., 2002; Faber et al., 2003; Goto and Lindoso, 2010), particularly in chronic skin lesions and mucosal lesions (Weigle et al., 1987; Weigle et al., 2002) as they harbour low parasite loads (Gutierrez et al., 1991; Jara et al., 2013). Furthermore, laboratory diagnosis of leishmaniasis remain challenging at health posts or centres that provide primary care in rural areas where TL is endemic due to the necessity of resources and infrastructure (Reimão et al., 2020). Molecular methods based on the polymerase chain reaction (PCR), including real-time PCR assays, are accessible only in research and reference laboratories and became increasingly important to complement the conventional parasitological tests. PCR-based methods provide the most sensitive and specific techniques used for *Leishmania* detection. Often, molecular typing of the infecting parasite species is necessary to guide the selection of the most appropriate treatment or prognosis of the disease (Arévalo et al., 2007; Van der Auwera and Dujardin, 2015; Akhoundi et al., 2017; Moreira et al., 2018; Mesa et al., 2020).

Recent research efforts have been devoted to develop and test molecular methods that have great potential to be further developed into point-of-care (POC) diagnostic tools for leishmaniasis and other neglected tropical diseases (reviewed in Bharadwaj et al., 2021). Among these, isothermal nucleic acid amplification techniques (NAATs) have attracted attention owing to their operation at a constant temperature (thus requiring minimal laboratory setup) and cost-effectiveness. Several studies that evaluated their diagnostic accuracy for different clinical forms of leishmaniasis reported high sensitivity (Adams et al., 2010; Mugasa et al., 2010; Saldarriaga et al., 2016; Adams et al., 2018; Ibarra-Meneses et al., 2018; Schallig et al., 2019; Cossio et al., 2021; Dixit et al., 2021; Travi et al., 2021). Still, these methods require further development and optimization for POC testing and field validation before their implementation in routine clinical care. Isothermal NAATs may suffer from non-specific amplification that can lead to low specificity, which can impact their applicability (Wang et al., 2015). Another test development concerns the simplified and standardized detection of PCR-amplified DNA of *Leishmania* using an oligochromatographic dipstick format (i.e., the commercially available *Leishmania* OligoC-TesT kit), which showed high diagnostic sensitivity and specificity in clinical samples (Deborggraeve et al., 2008; Espinosa et al., 2009). The advent of user-friendly portable devices such as Palm PCR™ (Kariyawasam et al., 2021), real-time fluorimeters [e.g. for real-time monitoring of the amplification profile of loop-mediated isothermal amplification (LAMP) reactions (Ibarra-Meneses et al., 2018; Dixit et al., 2021)], and the Bento Lab^®^, a mobile DNA laboratory setup (Kambouris et al., 2020), have opened the possibility to ease implementation of field-applicable molecular methods for use at POC to facilitate early diagnosis of leishmaniasis.

Most recently, the clustered regularly interspaced short palindromic repeats (CRISPR)/CRISPR-associated proteins (Cas) technology is revolutionizing the field of nucleic acid detection and next-generation molecular diagnostics, making possible field-deployable POC testing solutions (Myhrvold et al., 2018; Verosloff et al., 2021). CRISPR-Cas systems, naturally occurring in many bacteria and archaea as adaptive immune systems (Wiedenheft et al., 2012), are aiding in the development of novel nucleic acid detection platforms for major diseases such as cancer (Chen et al., 2018) and infectious diseases (Gootenberg et al., 2017). CRISPR-Cas technology is based on the specificity, programmability, and versatility of Cas effector proteins for nucleic acid sensing. Cas enzymes are easily programmable by the custom design of CRISPR RNAs (crRNAs) to recognize and cleave target DNA or RNA sequences with single-nucleotide specificity (Gootenberg et al., 2017). The most widely used Cas enzymes in nucleic acid detection and diagnostic applications are Cas9, Cas12a, and Cas13a. The latter two exhibit non-sequence-specific collateral (*trans*-cleavage) activity on target recognition. The discovery of the collateral cleavage activity on non-targeted single-stranded DNA (Cas12a) or single-stranded RNA (Cas13a) in solution funneled the development of CRISPR-Cas12a-based (DETECTR, Chen et al., 2018) and –Cas13a-based (SHERLOCK, Gootenberg et al., 2017) methods for *in vitro* nucleic acid detection. By adding quenched fluorescent oligonucleotide reporter probes in the CRISPR reaction mixture, which are cleaved by collateral activity, the released fluorescent signal indicates the presence of the target nucleic acid in a sample. Cas-mediated detection is commonly analyzed via fluorescence and lateral-flow readouts. Most CRISPR-based detection platforms function downstream of NAATs, which improves specificity due to crRNA-directed target sequence recognition (Kaminski et al., 2021). The preamplification step using NAATs enriches target molecules and increases the sensitivity of target detection down to attomolar (10^−18^ M) clinically meaningful concentrations (Gootenberg et al., 2017; Chen et al., 2018). Both isothermal (used in most CRISPR-based methods including SHERLOCK and DETECTR) and PCR-based (e.g. used in HOLMES combined with Cas12a-mediated detection, Li et al., 2018) NAATs are employed as preamplification strategies. CRISPR-based methods have been developed and applied to detect pathogens of public health importance, including bacteria [e.g. *Mycobacterium tuberculosis* (Ai et al., 2019; Sam et al., 2021)], viruses [e.g. Zika virus, dengue virus (Gootenberg et al., 2017; Gootenberg et al., 2018; Myhrvold et al., 2018), human papillomavirus (Chen et al., 2018; Gong et al., 2021), SARS-CoV-2 (Broughton et al., 2020; Fozouni et al., 2021; Alcántara et al., 2021a)], fungi (Huang et al., 2021), and protozoa [with a major focus on parasites belonging to the phylum Apicomplexa, such as *Plasmodium* spp. (Lee et al., 2020; Cunningham et al., 2021), *Toxoplasma gondii* (Ma et al., 2021), and *Cryptosporidium parvum* (Yu et al., 2021)]. Concerning trypanosomatid protozoan parasites, a recent study reported a newly developed CRISPR-dCas9-based DNA detection scheme with visual readout via DNAzymes (G-quadruplex-hemin complexes) and used spiked-in kinetoplast DNA (kDNA) from *Leishmania* into blood and urine samples as a proof-of-principle (Bengtson et al., 2022).

Here, we aimed to establish a novel method for CRISPR-Cas12a-mediated detection of amplified DNA from human lesion samples derived from patients with suspected cutaneous leishmaniasis (CL). Particularly, we used PCR amplification of multicopy *Leishmania* genetic targets coupled to downstream Cas12a-based detection aiming for high sensitivity and specificity of *Leishmania* detection with potential applicability in the diagnosis of CL in Latin America. To this end, we propose proof-of-concept of two assays, one targeting the conserved 18S ribosomal RNA gene (18S rDNA) for pan-*Leishmania* detection and the other targeting a region of kDNA minicircles conserved among *L*. (*Viannia*) species of medical importance in Latin America. We report the analytical sensitivity and specificity using extracted DNA from *Leishmania* and *Trypanosoma cruzi* reference strains. The assay performance was assessed using a panel of DNA samples from human skin lesion specimens with known diagnosis, wherein assay concordance was analyzed relative to a previously validated kDNA qPCR assay (Jara et al., 2013).

## MATERIALS AND METHODS

### Bioinformatics analyses

#### crRNA selection and template design

For the 18S rDNA gene, crRNA guide sequence candidates were obtained from EuPaGDT (http://grna.ctegd.uga.edu/; Peng and Tarleton, 2015) based on the *Leishmania* (*V*.) *braziliensis* MHOM/BR/75/M2904 sequence (TriTrypDB ID: LbrM.27.2.208540; at https://tritrypdb.org/tritrypdb/). The top ten protospacer adjacent motif (PAM) sequences (TTTV for LbCas12a, formerly LbCpf1) at the 5’ end of the target DNA and the 20nt-long target recognition sequences (i.e., guide sequence) in the crRNA were selected based on the total score and guide RNA efficiency/activity score. To identify conserved regions in *L*. (*Viannia*) kDNA minicircle sequences, an initial computational alignment included the sequences of 14 *L*. (*V*.) *braziliensis*, 10 *L*. (*V*.) *guyanensis*, 18 *L*. (*V*.) *lainsoni*, and 13 *L*. (*V*.) *panamensis* strains reported in GenBank. A second alignment focused on *L*. (*V*.) *braziliensis* kDNA minicircle sequences (409 sequences reported in GenBank as of June 4, 2020). All alignments were performed locally using the ClustalX package for Ubuntu with default parameters (Larkin et al., 2007). Conserved regions in the *Viannia* subgenus of *Leishmania* were identified with the AliView software (Larsson, 2014). PAM sequences were searched within the conserved regions to identify potential target recognition sites. Sequence alignments are available in **Supplementary File 1**.

Then, PAM sequences and recognition sites selected for both DNA targets were aligned against available sequences in GenBank of *L*. (*V*.) *braziliensis, L*. (*V*.) *guyanensis, L*. (*V*.) *panamensis, L*. (*V*.) *lainsoni*, and *L*. (*L*.) *major* using NCBI BLAST. For the 18S rDNA gene, sequences with less than 100% identity and coverage in any *Leishmania* species were discarded. For kDNA minicircles, sequences with less than 100% identity and coverage in any species from the *L*. (*Viannia*) subgenus were discarded.

In order to minimize the occurrence of cross-reactivity with the human genome, with microorganisms that cause skin lesions other than CL, and phylogenetically related protozoan parasites, filtered PAM sequences and target recognition sites were aligned against available genome sequences in GenBank of *Homo sapiens, Trypanosoma cruzi, Mycobacterium tuberculosis, Mycobacterium leprae, Sporothrix schenckii, Trypanosoma brucei, Blastomyces, Plasmodium vivax, Plasmodium falciparum*, and *Toxoplasma gondii*. Sequences were ranked by the number of different species of those listed above with at least 75% coincidence in sequence identity and coverage. For each target, the recognition site with the lowest rank was finally chosen.

In addition, a crRNA targeting the human RNase P *POP7* gene reported by Broughton et al. (2020) was used as a control for specimen quality. Double-stranded DNA (dsDNA) templates for crRNA generation through *in vitro* transcription were designed with a T7 promoter sequence (Beckert and Masquida, 2011; Alcántara et al., 2021b), followed by the LbCas12a crRNA scaffold (Chen et al., 2018) and the selected target recognition site.

#### Primer design

For the 18S rDNA gene, primer candidates for PCR preamplification of the target sequences prior to the CRISPR-Cas reaction were designed using the Primer3Plus v.2.4.2 server with the default settings and an average Tm of 60 °C (https://www.bioinformatics.nl/cgi-bin/primer3plus/primer3plus.cgi; Untergasser et al., 2012). For kDNA minicircles, primer candidates were manually searched (18 – 22nt long, average Tm of 60 °C) within conserved regions identified previously.

Primer candidates were aligned against the human genome using NCBI BLAST and discarded if they had more than 80% sequence identity and coverage with any human sequence. Self-and heterodimer formation were tested using the IDT OligoAnalyzer Tool (https://www.idtdna.com/pages/tools/oligoanalyzer). Primers with ΔG < -7 kcal/mol in any parameter were discarded. For the RNase P gene, primers reported previously (Curtis et al., 2018; Alcántara et al., 2021a) were selected. Oligonucleotides were ordered from Macrogen Inc. (Seoul, South Korea). All primer and crRNA template sequences used in this study are listed in **Table S1**.

### crRNA preparation

crRNAs were prepared from dsDNA templates using the TranscriptAid T7 High Yield Transcription Kit (Thermo Fisher Scientific, Waltham, MA, USA) at 37 °C for 3 h. Immediately thereafter, crRNAs were purified using the Direct-zol RNA miniprep kit (Zymo Research, Irvine, CA, USA) with a DNase I digestion in a column step for 15 min at room temperature (RT) and eluted in 80 μl nuclease-free water. Finally, crRNAs were quantified by the NanoDrop One microvolume UV-Vis spectrophotometer (Thermo Fisher Scientific), aliquoted to working volumes (∼ 10 μl) and stored at -80 °C (Alcántara et al., 2021b).

### DNA samples from reference strains of Leishmania spp. and Trypanosoma cruzi

Genomic DNA (gDNA) samples extracted from cultured promastigotes of reference strains of *Leishmania* [*L*. (*V*.) *braziliensis* (MHOM/BR/75/M2904, MHOM/PE/91/LC2043 and MHOM/PE/91/LC2177), *L*. (*V*.) *peruviana* (MHOM/PE/90/LCA08, MHOM/PE/90/HB22), *L*. (*V*.) *guyanensis* (IPRN/PE/87/Lp52), *L*. (*V*.) *lainsoni* (MHOM/BR/81/M6426), *L*. (*L*.) *amazonensis* LV79 (MPRO/BR/72/M1841), *L*. (*L*.) *major* (MHOM/SU/73/5-ASKH), *L*. (*L*.) *infantum* (MHOM/TN/80/IPT1), and *L*. (*L*.) *donovani* (MHOM/IN/80/DD8 (LEM 703)] analyzed here were retrieved from the DNA biobank of the leishmaniasis research group at the Molecular Epidemiology Unit of the Instituto de Medicina Tropical Alexander von Humboldt (IMTAvH), Universidad Peruana Cayetano Heredia (UPCH) in Lima, Peru. gDNA from the *Trypanosoma cruzi* Y strain (MHOM/BR/00/Y isolate; DTU TcII) was kindly provided by Dr. Manuela Verástegui (Infectious Diseases Research Laboratory, UPCH).

### Patient DNA samples

#### Ethics statement

This study used anonymized DNA samples derived from skin lesion specimens from patients who have tested positive or negative for *Leishmania* infection by conventional kDNA PCR (see below) and have given informed consent for research use of their specimens and clinical data. Patients with clinically suspected CL were recruited at the Hospital Nacional Adolfo Guevara Velasco (HNAGV) in Cusco, a region with endemic transmission of *Leishmania* (*Viannia*) parasites, with higher prevalence of *L*. (*V*.) *braziliensis* followed by *L*. (*V*.) *guyanensis* and *L*. (*V*.) *lainsoni* infections (Lucas et al., 1998; Sandoval-Juárez et al., 2020), during 2019 and 2020 as part of an ongoing collaborative study between the HNAGV and UPCH aimed at developing a rapid lateral flow assay for the diagnosis of TL in primary health care facilities in rural endemic areas in Peru. The protocol and informed consent of that study (registration number: 103155) were approved by the Institutional Review Board of the UPCH (IRB approval letter 063-05-19 dated 01/30/2019, renewed on 03/23/2021 with letter R-082-10-21). Written informed consent was obtained from all patients prior to enrollment. The activities of HNAGV and UPCH were conducted in compliance with all applicable international regulations governing the protection of human subjects.

#### Skin sample DNA isolation

DNA was isolated from different human skin lesion specimen types (biopsies, lancet scrapings, cytology brushes, swabs, and filter paper lesion impressions) as previously described (Boggild et al., 2010; Suárez et al., 2015). Samples were processed for DNA isolation using the High Pure PCR Template Preparation Kit (Roche, Mannheim, Germany) according to the manufacturer’s instructions. DNA from biopsy specimens was eluted into 150 μl of elution buffer, while for the other specimen types, a 100 μl elution volume was used. The isolated DNA was quantified by fluorometry using the Quant-iT high sensitivity dsDNA assay kit on the Qubit fluorometer (Invitrogen; Thermo Fisher Scientific). The DNA samples were stored at -20 °C until further molecular analyses.

### PCR-based preamplification of target DNA

PCR reactions were performed in a 25-μl reaction mixture consisting of 5 μl of DNA sample, 0.2 μM each target-specific forward and reverse primers (**Table S1**), and 1X DreamTaq Green PCR master mix (Thermo Scientific). Reactions were performed independently for kDNA, 18S rDNA, and RNase P. Cycling conditions consisted of an initial activation step at 95 °C for 2 min followed by 45 cycles of denaturation at 95 °C for 20 s, annealing at 60 °C (18S rDNA and RNase P) or 61 °C (kDNA) for 30 s, and extension at 68 °C for 15 s, followed by a final extension step at 68 °C for 5 min on the T100™ thermal cycler (Bio-Rad, Hercules, CA, USA). During protocol optimization, PCR products were visualized by 2%-3% agarose gel electrophoresis using SYBR Gold staining. A positive control (*L*. (*V*.) *braziliensis* M2904 gDNA, 10^4^ parasite genome equivalents/μl), a negative control (human gDNA from peripheral blood mononuclear cells (PBMC) of a healthy donor, 40 ng of input DNA), and two negative amplification controls (i.e. No-Template Control (NTC) reactions: NTC1, kept closed without water addition, and NTC2, made with PCR-grade water as the template replacement) were included in all experiments.

### LbCas12a trans-cleavage assays

LbCas12a-based detection reactions were performed as described previously (Chen et al., 2018; Broughton et al., 2020) with some modifications (Alcántara et al., 2021a). The recombinant LbCas12a protein was expressed and purified as described (Mendoza-Rojas et al., 2021). A single-stranded DNA (ssDNA) fluorophore quencher (FQ)-labeled reporter probe (5’ Cy3/TTATT/BHQ-2 3’) was selected and synthesized commercially (Macrogen Inc., Seoul, South Korea). The *in vitro* transcribed crRNA was first heated at 65 °C for 10 min in a heating block followed by refolding at RT for 10 min. Then, the CRISPR complex was prepared at 10-fold concentration (100 nM LbCas12a, 150 nM crRNA, 2 μM ssDNA-FQ reporter) in Reaction buffer (10 mM Tris-HCl (pH 7.9 at 25 °C), 50 mM NaCl, 100 μg/ml BSA, without adding MgCl2 at this step) and incubated at RT for 10 min in the dark. In parallel, 6 μl of PCR-amplified target DNA was diluted in 102 μl of Reaction buffer containing 18 mM MgCl2. Next, 10 μl of the CRISPR complex was mixed with 90 μl of the diluted PCR-amplified DNA in a flat-bottom, black 96-well microplate (Thermo Scientific; Cat. no. 237107). Final concentration of MgCl2 was 15 mM in the 100 μl final volume. Reactions were incubated in a fluorescence plate reader (most measurements used the Synergy™ H1 hybrid multi-mode reader, BioTek Instruments, Winooski, VT, USA; for experiments in **Figure 3, Supplementary Figures S2** and **S5**, measurements used the Cytation™ 5 Cell Imaging multi-mode reader, BioTek Instruments, see details in the respective figure legends) for 2 h at 25 °C. Fluorescence measurements were recorded every minute (excitation wavelength: 520 ± 9 nm, emission wavelength: 570 ± 20 nm) from the top of the wells. The fluorescence gain settings were 120 on the Synergy H1 plate reader and 150 on the Cytation 5 plate reader.

In addition to the PCR controls, a No-Template Control (NTC) of the CRISPR reaction (Reaction buffer containing 15 mM MgCl2 instead of the PCR product) was included. The PCR and CRISPR reaction setups were performed in a unidirectional workflow using separated laboratory work areas for each step to prevent amplicon carryover contamination.

### Analytical sensitivity and specificity testing

Serial dilutions of *L*. (*V*.) *braziliensis* M2904 gDNA (extracted from a promastigote culture) to a final range of 5 × 10^4^ to 5 × 10^−3^ parasite genome equivalents per reaction (encompassing the same range of parasite genome equivalents per reaction of the standard curve used in the kDNA qPCR assay; Jara et al., 2013) were tested as input DNA in PCR reactions to amplify the target gene, followed by LbCas12a-based detection assays as described above. The analytical sensitivity for each target gene was determined based on 3 independent experiments.

The analytical specificity of the PCR/CRISPR assays targeting *Leishmania* kDNA (*Viannia* subgenus) or 18S rDNA was tested using gDNA of laboratory reference strains of New World and Old World *Leishmania* species, and of *T. cruzi*. Target genes were amplified by PCR (using 20 – 40 ng of input DNA corresponding to 2.35 × 10^5^ – 4.71 × 10^5^ *Leishmania* genome equivalents) and detected by LbCas12a-based detection assays as described above. Two independent PCR/CRISPR experiments were performed.

### Performance evaluation of PCR/CRISPR assays on clinical samples

A total of 49 patient DNA samples extracted from skin lesion specimens described above were tested blindly in groups of 10, plus positive and negative controls. Five μl of 1/10 diluted DNA samples (1 – 250 ng of input DNA) were subjected to PCR-based amplification of *Leishmania* kDNA and 18S rDNA, and human RNase P gene. The PCR products were then detected by LbCas12a-based detection assays as described above. As a validation, a subset of samples was retested; this corresponded to 13 of 49 samples (26.5%) for kDNA PCR/CRISPR and 14 of 49 samples (28.6%) for 18S PCR/CRISPR.

### Conventional kDNA PCR

A qualitative conventional PCR targeting a 70-bp conserved region of *Leishmania* (*Viannia*) kDNA minicircles (López et al., 1993) was performed to detect *Leishmania* infection in clinical samples. The reaction mixture consisted of 0.4 μM of each primer (MP1L and MP3H; López et al., 1993) for amplification of the kDNA target, 0.4 μM of each primer (HBBL and HBBR; Boggild et al., 2010) for the amplification of the human beta-globin gene as an indicator of specimen adequacy, 0.2 mM dNTPs, 1X Qiagen PCR buffer, 1.5 mM MgCl2, 1X Q-Solution, 0.5 U HotStarTaq DNA polymerase (Qiagen, Hilden, Germany), and 5 μl of DNA sample (two reaction tubes were set up in parallel: one with the undiluted DNA sample and the other with 1/10 diluted DNA sample as a technical replicate) in a 25 μl total volume. Cycling conditions were as follows: 5 min at 95 °C followed by 40 cycles of 1 min at 94 °C, 1 min at 60 °C, 1 min at 72 °C followed by a final extension step for 10 min at 72 °C on the Veriti™ 96-well thermal cycler (Applied Biosystems; Thermo Fisher Scientific). Amplicons were visualized on 3% agarose gels stained with ethidium bromide. Each run included a positive control (DNA isolated from the *L*. (*V*.) *braziliensis* M2904 reference strain, 20 ng of input DNA), a negative control (human gDNA from PBMC of a healthy donor, 10-20 ng of input DNA), and a NTC.

### Quantitative real-time PCR

A SYBR Green-based qPCR assay targeting kDNA minicircles, based on the same primer set as for the diagnostic conventional PCR, was performed to detect and quantify *Leishmania* (*Viannia*) parasites in clinical samples, as described previously (Jara et al., 2013). A sample was considered detectable if a sigmoidal amplification curve and Cq value were obtained with a *Leishmania*-specific amplicon in the post-amplification melting curve analysis. The standard curve was prepared using 10-fold serially diluted *L*. (*V*.) *braziliensis* M2904 gDNA corresponding to 5 × 10^4^ to 5 × 10^−3^ parasite genome equivalents per reaction. In parallel, a qPCR assay detecting the human endogenous retrovirus 3 (*ERV-3*) gene, using the primers reported by Yuan et al. (2001), was used to quantify host cells, as described previously (Jara et al., 2013). The *ERV-3* standard curve was established from DNA extracted from human PBMC and comprised 2 × 10^4^ to 8 × 10^1^ copies/reaction. The *Leishmania* parasite load was calculated as follows: parasite genome equivalents (estimated by kDNA qPCR) normalized to the number of human cells (*ERV-3* average copy number/2) x 10^6^, expressed as the number of *Leishmania* parasites per 10^6^ human cells. Each run included a positive-control sample (DNA from a biopsy specimen of a patient with confirmed diagnosis of CL), which allowed monitoring the inter-assay variation in quantitative results between runs. In addition, a negative control (DNA from a non-leishmanial [kDNA PCR-negative] skin lesion) and a NTC were included. Standard curve dilution series, controls and clinical samples were tested in duplicate.

### Data processing and analysis

Following the LbCas12a-based detection reactions, the raw fluorescence collected data from each well of the assay plate were exported to Microsoft Excel. Based on the fluorescence time-course data gathered during the analytical sensitivity and specificity experiments with DNA samples from reference *Leishmania* strains and controls that were run in parallel, the time point of fluorescence accumulation for data analysis was defined (at 20-min time point for kDNA and at 10-min time point for 18S rDNA). For RNase P, analysis was performed at the 10-min time point as described previously (Broughton et al., 2020).

The raw fluorescence data were normalized by dividing target reaction fluorescence accumulated at the defined time point of the test sample to that of the NTC1 (PCR blank) reaction that was run in parallel on the same Cas12a assay plate. This is referred to as the ‘fluorescence ratio’. In the performance evaluation of the PCR/CRISPR assays with clinical samples, the fluorescence ratios were calculated using the mean of the NTC1 fluorescence values across all runs for a given target. In case of discordant results between the two independent measurements for the subset of clinical samples tested twice (i.e., discordant if one was positive and the other was negative), the sample was retested to confirm the result.

To determine the threshold cutoff value for detection of *Leishmania* in clinical samples by PCR/CRISPR, we calculated the mean and standard deviation of the fluorescence ratio of the negative clinical samples (n = 10). We considered positive signal in a reaction as being over 3 standard deviations higher than the mean fluorescence ratio of the negative samples. As an alternative method to set the detection threshold, we calculated the percentage positivity (PP) of the test samples relative to a positive control (*L*. (*V*.) *braziliensis* M2904 preamplified target DNA from 5 × 10^4^ parasite genome equivalents) that was run in each Cas12a assay plate. This method controls for inter-plate variability of the assay (Wright et al., 1993; Lejon et al., 2006; Zimic et al., 2009). The PP of the test samples was calculated as: PP = (raw fluorescence of the test sample/raw fluorescence of the positive control of the corresponding plate) × 100. Then, the mean PP of the two independent measurements for the subset of retested samples was calculated. Using the PP of the tested samples as the predictor variable, a simple logistic regression was performed to model the *Leishmania* infection status (positive/negative, as determined by kDNA qPCR). Sensitivity, specificity and a receiver operating characteristic (ROC) curve were calculated over the range of cutoff points for the predictor variable (PP). The optimal probability cutoff point for classification (Pr-cutoff/PP) was selected based on the maximization of the Youden’s J-index (sensitivity + specificity – 1).

As quality control during CRISPR-Cas assay data analysis, we calculated the mean and standard deviation of the raw fluorescence values of the negative controls examined along with clinical samples across all runs at the 20-, 40-, 60- and 120-min time points. We established the range of acceptable fluorescence values for the negative controls to those that fell within 15% of the mean fluorescence (for each negative control) across all runs. In cases where any of the negative controls was out of the acceptable range, i.e. showing a marked increase in raw fluorescence, the experimental run was considered invalid due to contamination.

The CRISPR-Cas assay data analysis was blinded to the qPCR data and only compared once all samples had been tested. The concordance of the results obtained with kDNA PCR/CRISPR, 18S PCR/CRISPR, or kDNA conventional PCR (routine diagnostic test; López et al., 1993) against the kDNA qPCR reference method was assessed by calculating the positive and negative percent agreement (PPA and NPA, respectively) with Wilson score 95% confidence intervals (CI) using the Analyse-it software add-in for Microsoft Excel (https://analyse-it.com/). In addition, Cohen’s Kappa coefficient was calculated using kDNA qPCR as the reference method (McHugh, 2012).

Graphs, numerical data analyses, and statistical analyses were performed using GraphPad Prism version 9 (GraphPad Software, San Diego, CA, USA). For comparison of the fluorescence ratio data between groups of *L*. (*Viannia*) and *L*. (*Leishmania*) strains of the Cas12a detection experiments in **Figure 4**, an unpaired *t*-test was used. Pairwise comparisons of the fluorescence ratio data of reactions containing *Leishmania* target DNA (positive control) or human gDNA (negative control) with the no-template control (NTC2-PCR) of the Cas12a detection experiments analyzed in **Figure 8** were done using an unpaired *t*-test. Differences with *P* < 0.05 were considered significant. The logistic regression and ROC curve analysis were conducted using Stata version MP 17 (StataCorp, College Station, TX, USA) as well as GraphPad Prism version 9.

## RESULTS

### Development, optimization and analytical validation of Cas12a-based assays for detection of Leishmania spp

Herein, we sought to develop CRISPR-Cas12a-based assays for genus-specific detection of *Leishmania* spp. and *L*. (*Viannia*) subgenus-specific detection of the species responsible for most cases of cutaneous and mucosal leishmaniasis in Latin America. We chose the widely used multicopy 18S rDNA and kDNA minicircle targets to ensure high sensitivity and to provide proof-of-concept of the applicability of CRISPR-based detection to *Leishmania* spp. in clinical specimens. Each assay starts from extracted DNA and comprises a preamplification step of the target DNA using conventional PCR, followed by Cas12a detection with fluorescent readout on a plate reader (**Figure 1**). Thus, this test yields qualitative detection of nucleic acids from *Leishmania*. Raw fluorescence values over 2 h were measured to assess the fluorescence change over time for the test samples as compared to the controls. For assay interpretation and reporting results, the fluorescent signal of each sample obtained at a defined time point (20 min for kDNA and 10 min for 18S rDNA; see details below) of the Cas12a reaction was normalized relative to the No-Template Control (NTC) fluorescence value, thereby resulting in a fluorescence ratio.

**Figure 1.**
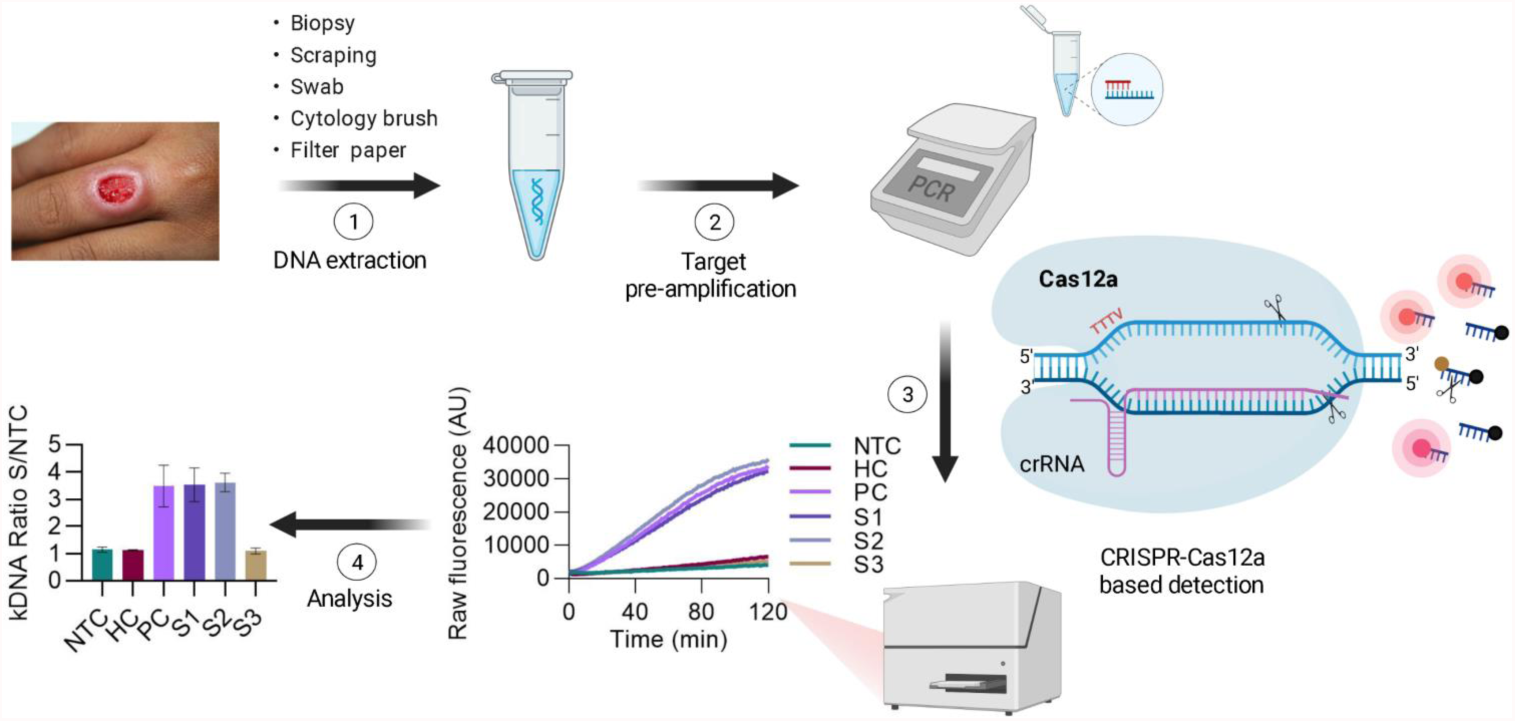
Schematic of the PCR coupled CRISPR-Cas12a-based assay workflow for detection of *Leishmania* spp. in clinical samples. (**1**) Skin lesion specimens suggestive of CL are processed for gDNA extraction. (**2**) The isolated DNA is subjected to PCR preamplification using primers targeting *Leishmania* kDNA minicircles or 18S rDNA, or the sample control, human RNase P gene. (**3**) The PCR amplicon is then used as input DNA for Cas12a-based detection guided by a target-specific crRNA. A ternary complex is formed if the target DNA is present in the reaction mixture. Cas12a recognizes a 5’-TTTV-3’ PAM in the DNA target, which results in base pairing between the crRNA guide (spacer) segment and the complementary target DNA. Upon specific target DNA recognition by Cas12a, the activated Cas12a enzyme generates non-sequence-specific cleavage of a single-stranded DNA (ssDNA) reporter probe, thereby producing a fluorescence signal that can be recorded by a fluorescence plate reader. (**4**) Normalized fluorescence readings (test sample/NTC), expressed as a fluorescence ratio, are used to interpret the assay results. As an illustrative example, data shown correspond to the kDNA target and are represented as mean ± standard deviation (SD) (n = 2 independent amplification and detection runs). Standardized conditions consisted of ∼90-minute amplification (45 PCR cycles) and fluorescent signal (Cas12a assay) at 20-min time point for kDNA and at 10-min time point for both 18S rDNA and RNase P. AU, arbitrary units; NTC, no-template control of the Cas12a reaction; HC, human PBMC gDNA from a healthy donor (negative control); PC, *L*. (*V*.) *braziliensis* M2904 gDNA (positive control); S1-S3, DNA from clinical samples. Figure created with BioRender.com.

The highly conserved nature of the 18S rDNA gene was used for the development of a pan-*Leishmania* detection assay using a crRNA directed to a conserved target site across *Leishmania* species (**Figures 2A,B** and **Supplementary File 1**). For the kDNA target, we first searched *in silico* for conserved regions in kDNA minicircle sequences among New World *L*. (*Viannia*) species and selected one region with a suitable PAM site for Cas12a crRNA design (**Figures 2A,B, Supplementary File 1**, and **Supplementary Figure S1**). We designed a kDNA-crRNA, located within minicircle conserved sequence block CSB2 (**Supplementary Figure S1**), which recognizes a target sequence that is highly conserved among *L*. (*Viannia*) species according to the *in silico* multiple sequence alignments performed (**Figure 2B** and **Supplementary File 1**). A careful crRNA selection was carried out for both *Leishmania* target sequences, thereby filtering against the human genome and genomes from related pathogens that co-circulate in leishmaniasis endemic regions and/or cause leishmaniasis-like lesions (**Figure 2A**).

**Figure 2.**
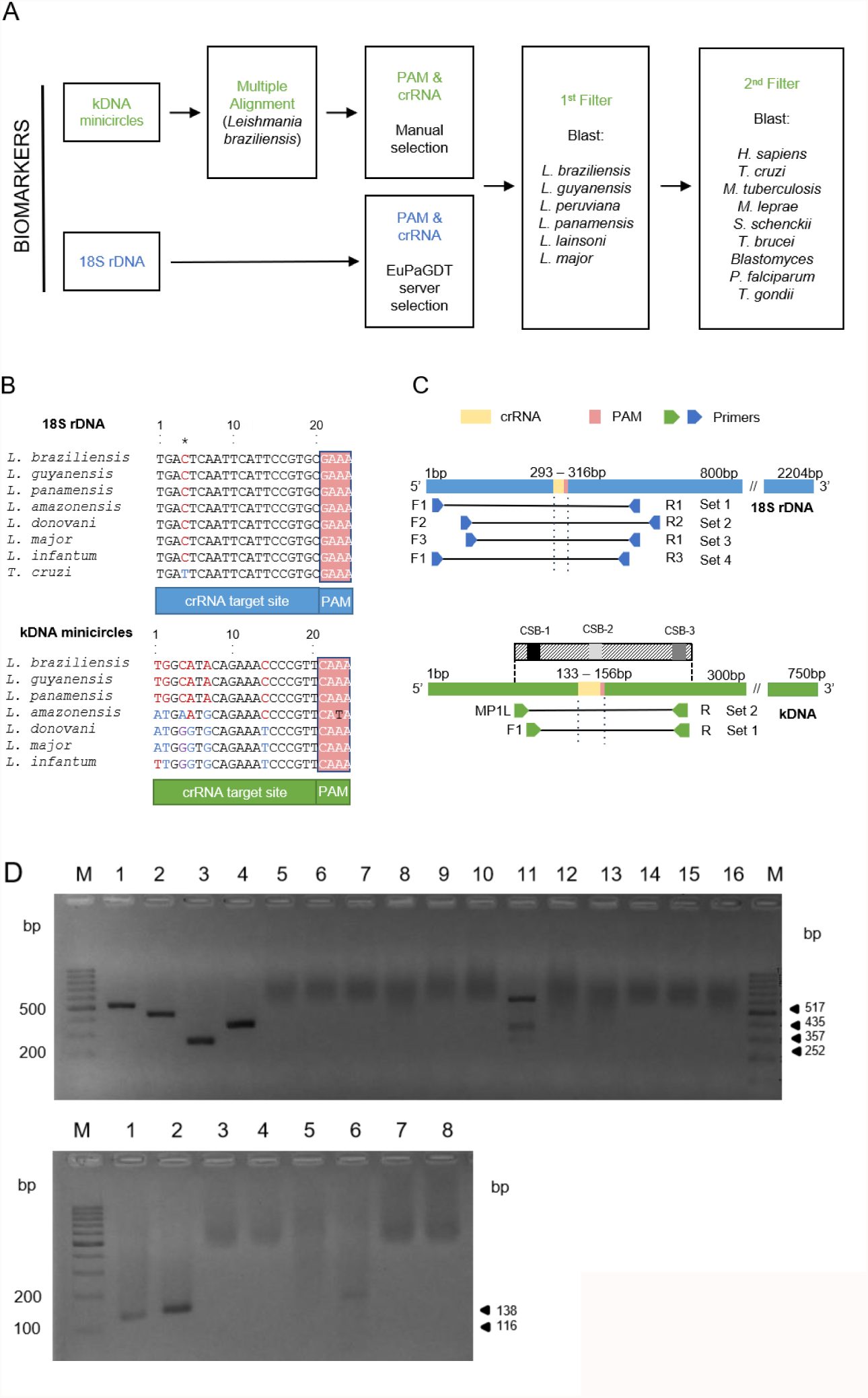
Bioinformatics analysis for crRNA and primer sets design. (**A**) Bioinformatics workflow for crRNA filtered selection for *Leishmania* kDNA minicircle and 18S rDNA biomarkers (see Materials and Methods for details). (**B**) Alignments of the target DNA sequence (that is complementary to the crRNA guide sequence) and flanking PAM from 7 strains representative of different *Leishmania* species for the two targets and the *T. cruzi* Y strain for the 18S rDNA target (the kDNA alignment including *T. cruzi* Y strain resulted in a 21 bp gap in the crRNA). Aligned sequences of the 18S rDNA gene concerned the strains: *L*. (*V*.) *braziliensis* MHOM/BR/75/M2904 2019 (TriTrypDB ID code: LbrM.27.2.208560), *L*. (*V*.) *guyanensis* MHOM/BR/1975/M4147 (GenBank accession number: X53913), *L*. (*V*.) *panamensis* MHOM/PA/94/PSC-1 (TriTrypDB ID code: LPMP_27rRNA1), *L*. (*L*.) *amazonensis* MHOM/BR/1973/M2269 (TriTrypDB ID code: LAMA_000552800), *L*. (*L*.) *donovani* MHOM/ET/67/HU3 (isolate LV9, TriTrypDB ID code: LdLV9.00.2.200020), *L*. (*L*.) *major* MHOM/IL/80/Friedlin (TriTrypDB ID code: LmjF.27.rRNA.06), *L*. (*L*.) *infantum* MCAN/ES/98/LLM-877 (strain JPCM5, TriTrypDB ID code: LINF_270031400), and *T. cruzi* Y (GenBank accession number: AF301912). The alignment of the nucleotide sequences of the kDNA minicircle target shown here included the strains (GenBank accession numbers are indicated between brackets): *L*. (*V*.) *braziliensis* MHOM/BR/75/M2904 (KY698821), *L*. (*V*.) *guyanensis* MHOM/BR/78/M5378 (KY699068), *L*. (*V*.) *panamensis* MHOM/PA/75/M4037 (AF118474), *L*. (*L*.) *amazonensis* RAT/BR/72/LV78 (KY698896), *L*. (*L*.) *donovani* MHOM/IN/80/DD8 (AF167712), *L*. (*L*.) *major* MHOM/IL/67/LV563 (KM555295), *L*. (*L*.) *infantum* MHOM/FR/91/LEM-2298 (AF190883). Nucleotide positions differing among aligned sequences are highlighted. The 18S rDNA target sequence shows conservation at the *Leishmania* genus level, whereas the selected kDNA minicircle target region is conserved among *L*. (*Viannia*) species. See Supplementary File 1 for all alignments. (**C**) Locations of PCR primer sets that flank the crRNA target site and PAM sequence in the target genomic regions (indicated nucleotide positions are based on the *L*. (*V*.) *braziliensis* M2904 strain). For the kDNA minicircle target, we used PCR primers located within conserved sites CSB1 and CSB3 (Supplementary Figure S1). (**D**) Agarose gel electrophoresis of PCR products generated with each of the six primer sets shown in panel **C**. M, molecular size marker (100 bp; GeneRuler, Thermo Scientific). The PCR amplicons (6 μl) were mixed with 2 μl 6X TriTrack DNA loading dye (Thermo Scientific) plus SYBR Gold solution, loaded onto a 2% agarose gel (for 18S rDNA) or 3% agarose gel (for kDNA), and run in 1X TBE buffer for 1 hour at 100V. *Upper gel*: PCR products obtained for the amplified region of the 18S rDNA target using the 18S primer set 1 (lanes 1, 5, 9, 13), set 2 (lanes 2, 6, 10, 14), set 3 (lanes 3, 7, 11, 15), and set 4 (lanes 4, 8, 12, 16). Lanes 1–4: *L*. (*V*.) *braziliensis* M2904 gDNA (5 × 10^4^ parasite genome equivalents per PCR reaction; positive control). The size of the expected product is 517 bp for primer set 1 (lane 1), 435 bp for primer set 2 (lane 2), 252 bp for primer set 3 (lane 3), and 357 bp for primer set 4 (lane 4). Lanes 5–8: NTC1 (kept closed without water addition). Lanes 9–12: human PBMC gDNA (HC, negative control). The non-specific bands seen in the HC reaction with the 18S primer set 3 (lane 11) did not interfere with the Cas12a assay (see Figures 3B and 4B). Lanes 13–16: NTC2 (made with water instead of the template). *Bottom gel*: PCR products obtained for the amplified region of the kDNA target using the kDNA primer set 1 (lanes 1, 3, 5, 7) and set 2 (lanes 2, 4, 6, 8). Lanes 1 and 2, *L*. (*V*.) *braziliensis* M2904 gDNA (5 × 10^4^ parasite genome equivalents per PCR reaction; positive control). The size of the expected product is 116 bp for primer set 1 (lane 1) and 138 bp for primer set 2 (lane 2). Lanes 3 and 4, NTC1. Lanes 5 and 6, HC (negative control). The non-specific band seen in the HC reaction with the kDNA primer set 2 (lane 6) did not interfere with the Cas12a assay (see Supplementary Figures S2A-D). Lanes 7 and 8, NTC2.

We then designed PCR primers to amplify genomic segments spanning the selected crRNA target sites (**Figure 2C** and **Table S1**). For the 18S rDNA target, we designed three forward and three reverse primers to be tested in four primer pairs (sets 1–4, **Figure 2C**). For the kDNA target, we designed PCR primers located within minicircle conserved sequence blocks CSB1 and CSB3 (kDNA primer set 1, namely primers F1/R, **Figure 2C** and **Supplementary Figure S1**). Additionally, we selected a combination of the previously published MP1L primer (López et al., 1993), located immediately upstream of the CSB1, and the designed R primer located within CSB3 (kDNA primer set 2, MP1L/R, **Figure 2C** and **Supplementary Figure S1**).

During the optimization of PCR amplification conditions for both *Leishmania* targets, each primer set generated a PCR product of the expected size from genomic DNA of the *L. braziliensis* M2904 reference strain, while not producing an amplified product from the NTC controls (**Figure 2D**). The standardized conditions consisted of ∼90-minute amplification (45 PCR cycles), in order to boost the sensitivity of the PCR/CRISPR assays. Two primer pairs (kDNA primer set 2 and 18S primer set 3) produced non-specific PCR products in the negative control with human genomic DNA (**Figure 2D**). However, these did not generate a signal above the NTC controls in the Cas12a-based detection assay (see below).

For our Cas12a-based assays we used conditions that were previously optimized for viral targets of SARS-CoV-2 (Alcántara et al., 2021a; Alcántara et al., 2021b), namely 10 nM LbCas12a, 15 nM crRNA, 15 mM MgCl2, and 200 nM of the reporter probe. The Cas12a reaction was incubated at 25 °C as previously described (Alcántara et al., 2021a; Alcántara et al., 2021b). Next, we carried out serial dilutions of *L. braziliensis* M2904 genomic DNA used as template DNA to determine the analytical sensitivity of the PCR-based preamplification with each primer set coupled to Cas12a-based detection. The raw fluorescence curves over 2 h clearly delineated a positive result (i.e., target DNA detected) from a negative one or background with a marked increase in relative fluorescence for both *Leishmania* kDNA (**Figure 3A**) and 18S rDNA (**Figure 3B**) targets. For the kDNA target, on the basis of the raw fluorescence curves (**Figure 3A**) a time point 20 min discriminated between specific and background signal from the human negative control (HC) and NTC controls. Preamplification of the kDNA target with either primer set 1 or set 2 allowed detection of at least 5 × 10^−2^ parasite genome equivalents/reaction (**Figures 3C,D**). Moreover, in one out of 3 independent experiments using kDNA primer set 2, detection of 5 × 10^−3^ parasite genome equivalents/reaction (fluorescence ratio of the Cas12a assay = 3.43) was achieved (**Figure 3D**). Thus, we found mostly similar analytical sensitivity of the kDNA PCR/CRISPR assay using either kDNA primer set 1 or set 2 coupled to crRNA-guided Cas12a detection as compared to the performance of the kDNA qPCR assay for *Leishmania* DNA detection (Jara et al., 2013).

**Figure 3.**
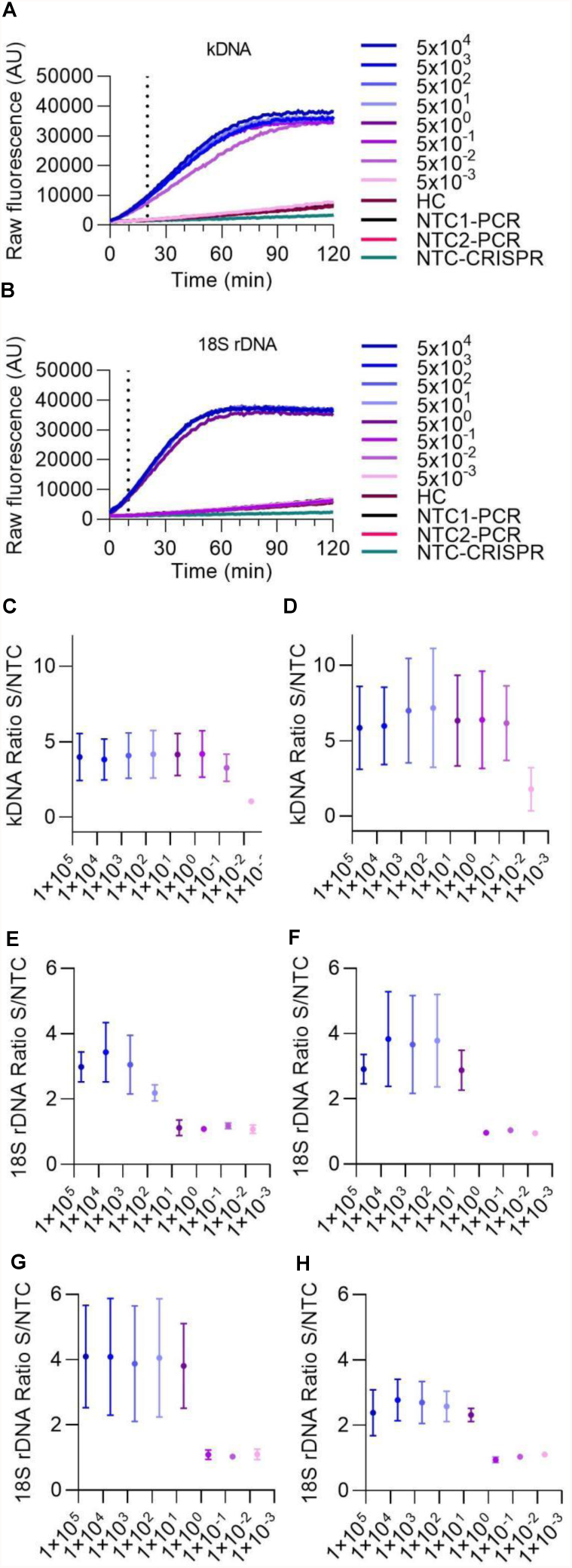
Analytical sensitivity of Cas12a-based detection of *L*. (*V*.) *braziliensis* M2904 DNA. Eight serial dilutions of *L*. (*V*.) *braziliensis* M2904 gDNA were subjected to PCR amplification followed by Cas12a-based detection for kDNA and 18S rDNA targets, for determination of the lowest amount of parasite genome equivalents that can be consistently detected. Different primer sets (shown in Figure 2C) were evaluated to determine those with the best performance. (**A**,**B**) Raw fluorescence signal from the Cas12a reaction over 2 h on the amplicon generated with the kDNA primer set 1 (**A**) or 18S primer set 3 (**B**). Data from one experiment are shown, with fluorescence measurements taken on the Synergy H1 plate reader. (**C**,**D**) Data points depict the fluorescence ratio (fluorescence signal obtained at 20-min time point in the test sample relative to the NTC) from Cas12a reactions on the amplicon generated with the kDNA primer set 1 (**C**) or set 2 (**D**). Data are represented as mean ± SD (n = 3 independent amplification and detection runs). The X axis has a logarithmic scale. (**E-H**) Data points depict the fluorescence ratio (fluorescence signal obtained at 10-min time point in the test sample relative to the NTC) from Cas12a reactions on the amplicon generated with the 18S primer set 1 (**E**), set 2 (**F**), set 3 (**G**), or set 4 (**H**). Data are represented as mean ± SD (n = 3 independent amplification and detection runs). The X axis has a logarithmic scale. In **C**,**D** and **F**, fluorescence measurements of Cas12a detection runs were taken on the Synergy H1 plate reader (2 runs) and on the Cytation 5 plate reader (one run). In **E** and **G**, one detection run was performed on the Synergy H1 plate reader and 2 detection runs were made on the Cytation 5 plate reader. In **H**, all 3 detection runs were conducted on the Cytation 5 plate reader.

For the 18S rDNA target, the fluorescence signal of the Cas12a detection assay showed saturation in less than 60 min (**Figure 3B**); and a time point 10 min allowed us to distinguish between specific and background signal. The Cas12a assay employing preamplified 18S rDNA target with either primer set 2 (**Figure 3F**), set 3 (**Figure 3G**) or set 4 (**Figure 3H**) could detect at least 5 × 10^0^ parasite genome equivalents/reaction, whereas target preamplification with primer set 1 (**Figure 3E**) resulted in lower detection sensitivity, i.e. of at least 5 × 10^1^ parasite genome equivalents/reaction. Of these evaluated 18S primer sets, we chose primer set 3 out of convenience for further evaluation.

We then evaluated the specificity of our PCR/CRISPR assays with DNA samples from 11 reference strains of various *L*. (*Viannia*) and *L*. (*Leishmania*) species as well as one strain of the phylogenetically closely related *T. cruzi* (**Figure 4**). For the PCR-based preamplification step, we tested both kDNA primer sets and the 18S primer set 3. For the kDNA target, the crRNA-guided Cas12a detection assay on preamplified target DNA with primer set 1 detected a distinctive fluorescent signal with *L*. (*Viannia*) DNA [fluorescence ratio values at 20-min time point ranging from 1.816 to 2.817; n = 7 strains], whereas DNA samples from representative strains of *L*. (*Leishmania*) species [range, 0.827 – 1.229; n = 4 strains] and the *T. cruzi* Y strain [range, 0.892 – 0.915] did not exhibit signal above the negative control reactions [range, 0.864 – 1.179] (**Figure 4A** and associated time course data in **Supplementary Figures S3A-D**). In contrast, the Cas12a assay on the kDNA target preamplified with primer set 2 detected a comparable signal with any of *Leishmania* DNAs tested from strains belonging to the *L*. (*Viannia*) and *L*. (*Leishmania*) subgenera, while no cross-reaction was observed with *T. cruzi* Y or human DNA (**Supplementary Figures S2A-D**). Since we were interested in developing a *L*. (*Viannia*)-specific detection assay targeting kDNA minicircles, we selected kDNA primer set 1 for the preamplification step in combination with the examined kDNA-crRNA for subsequent testing of patient samples. For the 18S target, the Cas12a assay employing preamplified target DNA with primer set 3 detected a prominent fluorescent signal with DNA from all tested *Leishmania* species [fluorescence ratio values at 10-min time point ranging from 4.350 to 5.723; n = 11 strains], whereas the signal from the *T. cruzi* Y strain DNA [range, 0.885 – 0.962] could not be distinguished from the negative controls [range, 0.922 – 1.242] (**Figure 4B** and associated time course data in **Supplementary Figures S3E-H**). Therefore, the 18S primer set 3 was further used for the target preamplification step, in conjunction with the designed 18S-crRNA, in the evaluation of clinical samples.

**Figure 4.**
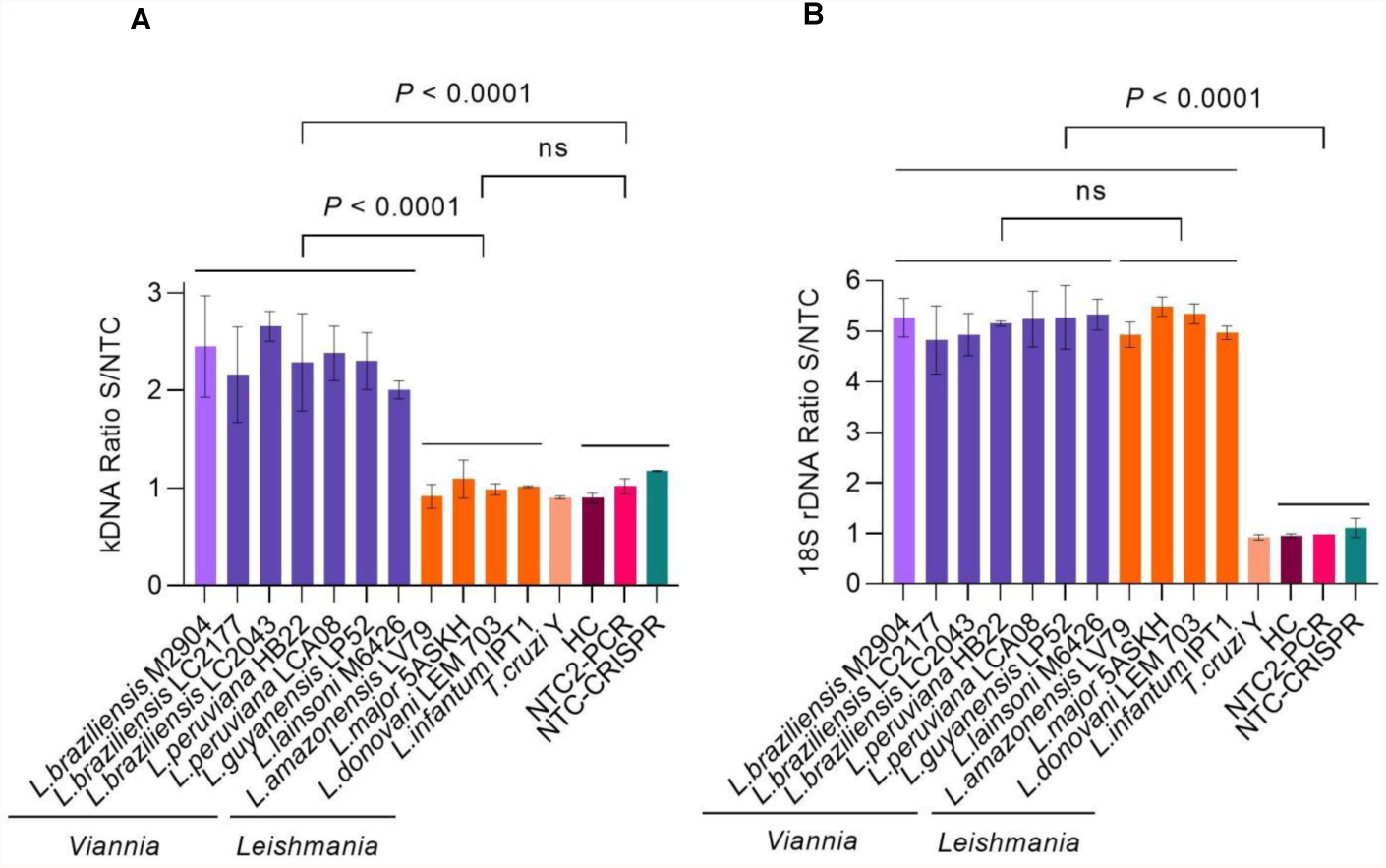
Analytical specificity of Cas12a-based detection of *Leishmania* spp. using reference strains. (**A**) The specificity of the kDNA PCR/CRISPR assay was evaluated using gDNA (preamplified with kDNA primer set 1) from 11 *Leishmania* strains (7 of subgenus *Viannia* and 4 of subgenus *Leishmania*) and the *T. cruzi* Y strain. (**B**) The same DNA samples from reference strains as in **A** were tested in the specificity assessment of the 18S PCR/CRISPR assay using the 18S primer set 3 in the preamplification step. Bar graphs depict the fluorescence ratio (fluorescence signal obtained at 20-min time point for kDNA and at 10-min time point for 18S rDNA in the test sample relative to the NTC) from Cas12a reactions. Data are represented as mean ± SD (n = 2 independent amplification and detection runs). Statistical comparisons between groups were conducted using an unpaired *t*-test (two-tailed *P* values are shown; non-significant *P* values are indicated by ns). Negative controls included human PBMC gDNA and NTC controls of PCR and CRISPR reactions. Fluorescence measurements of Cas12a detection runs were taken on the Synergy H1 plate reader. Full time course data are provided in Supplementary Figure S3.

### Performance evaluation of PCR/CRISPR assays in clinical samples

We next assessed the applicability of our CRISPR-based assays in clinical samples. We tested extracted DNA from 49 cutaneous lesion specimens from patients with previously confirmed positive (36/49; 73.5%) or negative (13/49; 26.5%) *Leishmania* infection status using the routine kDNA conventional PCR (cPCR) diagnostic test (**Figure 5A**). These samples were analyzed by the newly developed PCR/CRISPR assays targeting *Leishmania* kDNA and the 18S rDNA gene, as well as the human RNase P gene as a sample control, in parallel with a previously validated kDNA qPCR test (**Figure 5A**). Of the 49 samples analyzed, the kDNA qPCR test identified 79.6% of samples (39 of 49) as *Leishmania* positive, whereas 20.4% of samples (10 of 49) were negative.

**Figure 5.**
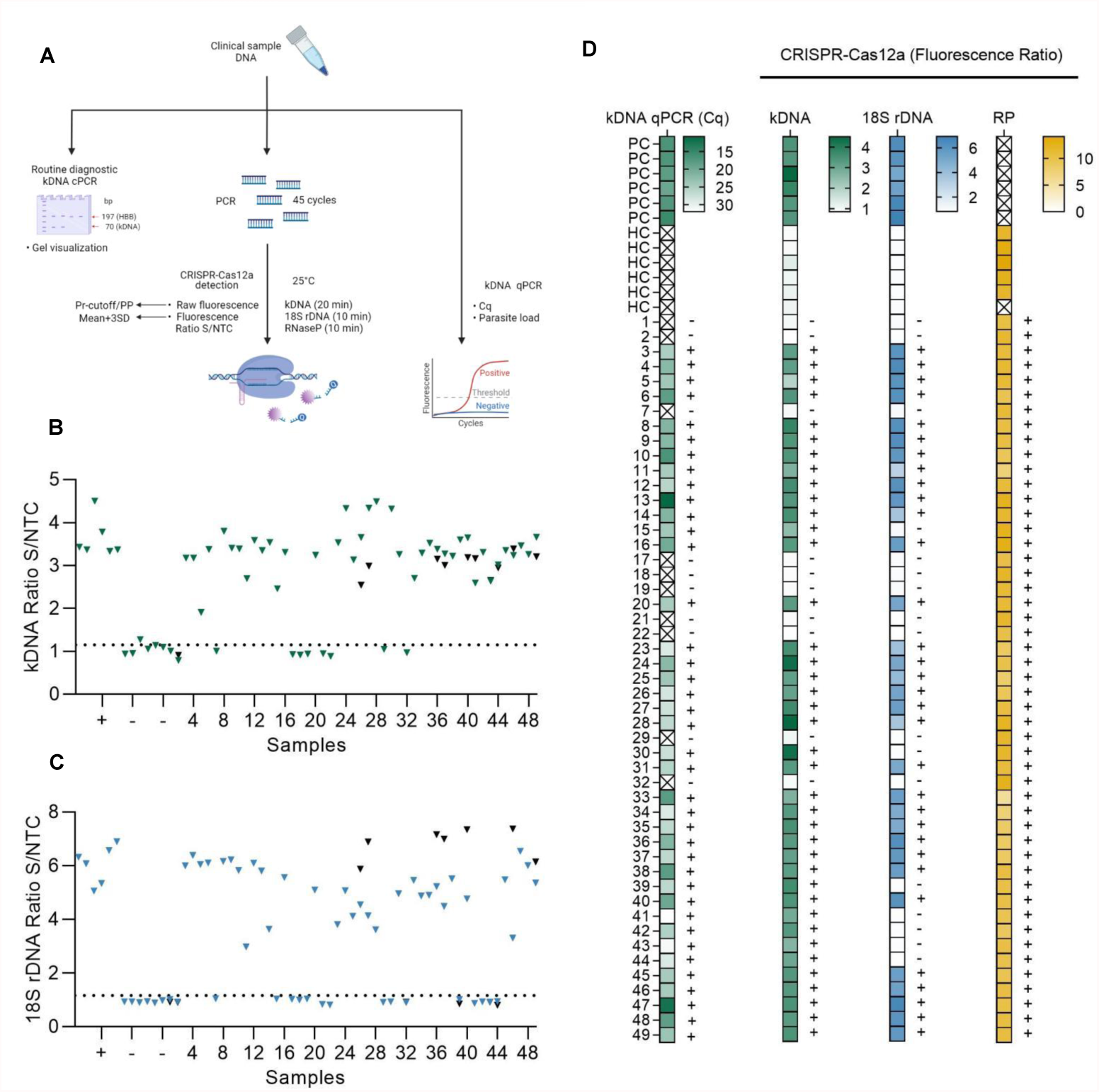
Performance evaluation of PCR/CRISPR assays for detection of *Leishmania* DNA targets in clinical samples. (**A**) Sample analysis workflow including molecular analyses (kDNA cPCR, kDNA qPCR, and PCR/CRISPR targeting *Leishmania* kDNA and 18S rDNA or human RNase P gene) and data analysis (see Materials and Methods for more details). Figure created with BioRender.com. (**B**,**C**) Extracted DNA from 49 clinical samples was subjected to PCR amplification using primers specific to kDNA (**B**) and 18S rDNA (**C**) of *Leishmania*, followed by Cas12a-based detection. Fluorescence measurements were made on the Synergy H1 plate reader. Data shown represent the fluorescence ratio (fluorescence signal obtained at 20-min time point for kDNA and at 10-min time point for 18S rDNA in the test sample relative to the NTC) from Cas12a reactions. The dashed lines indicate the positive threshold cutoff value (mean of the fluorescence ratio of negative clinical samples + 3SD; n = 10) for detection of *Leishmania* targets (cutoff value = 1.151 for kDNA and 1.171 for 18S rDNA). +, positive control (*L*. (*V*.) *braziliensis* M2904 gDNA; n = 6); -, negative control (human DNA from PBMC of a healthy donor; n = 6). A subset of clinical samples was retested (black symbols). Full time course data for a group of clinical samples are provided in Supplementary Figure S4. (**D**) Heat maps displaying qPCR and PCR/CRISPR data from clinical samples. *Left panel*, the color scale represents Cq values determined by kDNA qPCR. The X mark inside the box symbol indicates that no Cq value was obtained. *Second-fourth panels*, the color scale represents fluorescence ratio values from Cas12a reactions for *Leishmania* kDNA and 18S rDNA targets and the sample control, human RNase P (RP) gene, respectively. The X mark inside the box symbol in the RP gene heatmap indicates not applicable or not done. The detection threshold of the Cas12a assay on the *Leishmania* target was set as the mean of the fluorescence ratio of negative clinical samples (n = 10) + 3SD. The positive threshold for the RP gene was set as a fluorescence signal of 5-fold above background (i.e., fluorescence ratio > 5; Khan et al., 2021). The test results (+, detected; -, not detected) are indicated on the right side of the heat maps. Samples are coded as 1 to 49. PC, positive control (*L*. (*V*.) *braziliensis* M2904 gDNA for the PCR/CRISPR assay; DNA from a biopsy specimen positive by kDNA cPCR used for the qPCR assay). HC, human gDNA (from PBMC of a healthy donor). See Supplementary File 2 for the complete dataset of this study.

Considering the kDNA qPCR assay as the reference test for *Leishmania* detection, 39 of the 39 qPCR positive samples were also positive with the kDNA PCR/CRISPR assay using a cutoff equal to the mean + 3SD of the fluorescence ratio of 10 negative clinical samples (cutoff = 1.151). Fluorescence ratio values of positive samples by kDNA PCR/CRISPR ranged from 1.91 to 4.49 (**Figures 5B, 5D** and **Supplementary File 2**). Clinical samples that scored positive showed robust fluorescence curves from Cas12a detection of *L*. (*Viannia*) kDNA minicircle molecules (**Supplementary Figures S4A**,**B**). Repeated measurements for a subset of clinical samples indicated overall consistent results (**Figure 5B** and **Supplementary File 2**). However, two samples (CL-01 and CL-32) out of 13 retested samples showed discordant results (one of two replicates with positive signal in the kDNA Cas12a assay) but were suspected to be negative because both the kDNA cPCR and kDNA qPCR tests identified these samples as negative. To clarify the results, we repeated testing of these samples by kDNA PCR/CRISPR in duplicate, where both samples showed undetectable results (two of two replicates) (**Supplementary Figures S5A-D**).

The 18S PCR/CRISPR assay resulted in 32 of the 39 qPCR positive samples being classified as *Leishmania* positive (using a classification cutoff value of 1.171), with fluorescence ratio values ranging from 2.98 to 7.37 (**Figures 5C,D** and **Supplementary File 2**). Positive patient samples exhibited strong fluorescence curves in the Cas12a assay indicating presence of the *Leishmania* 18S rDNA gene (**Supplementary Figures S4C**,**D**). Consistent results were obtained upon repeated measuring on a subset of clinical samples (**Figure 5C** and **Supplementary File 2**). Discordance between test results (one of two replicates with positive signal in the 18S Cas12a assay) occurred in one sample (CL-41) out of 14 retested samples. That sample had high Cq values (mean Cq of 31.91) (**Figure 5D**) and low parasite load (4.61 parasites per 10^6^ human cells) (**Supplementary File 2**) as determined by the kDNA qPCR test. We repeated testing of sample CL-41 by 18S PCR/CRISPR in duplicate, confirming it as negative (two of two replicates) (**Supplementary Figures S5E**,**F**).

The human RNase P gene was detected in all 49 clinical samples analyzed by PCR/CRISPR (**Figure 5D** and **Supplementary File 2**), thereby confirming the quality of the samples and validating the process of DNA extraction from clinical specimens. Robust fluorescence curves were obtained for the RNase P gene from Cas12a detection (**Supplementary Figures S4E**,**F**), with fluorescence ratio values > 5 in all cases (**Supplementary File 2**).

Patient samples detected as *Leishmania* positive in the kDNA and 18S PCR/CRISPR assays showed a large range of Cq values (**Figure 6A** and **Supplementary File 2**) and, accordingly, covered a wide range of parasite loads (**Figure 6B** and **Supplementary File 2**) as determined by kDNA qPCR. The kDNA PCR/CRISPR assay detected the presence of *Leishmania* kDNA molecules in samples with parasite equivalents/reaction as low as 2.9 ×10^−3^ (sample CL-43) corresponding to a parasite load of 1 parasite per 10^6^ human cells (**Figure 6B** and **Supplementary File 2**). The *Leishmania* 18S rDNA target was detected by PCR/CRISPR in samples with parasite equivalents/reaction as low as 5.6 × 10^−2^ (sample CL-23) corresponding to a parasite load of 42 parasites per 10^6^ human cells (**Figure 6B** and **Supplementary File 2**). However, not all samples with parasite loads in the range between 35 and 100 parasites per 10^6^ human cells were detected by the 18S CRISPR-based assay (**Figure 6B** and **Supplementary File 2**). The 18S rDNA gene was consistently detected in samples containing parasite equivalents/reaction of at least 1 × 10^0^ corresponding to a parasite load greater than 10^2^ parasites per 10^6^ human cells (**Figure 6B** and **Supplementary File 2**). As expected, an inverse linear correlation between Cq values and quantified parasite load levels in clinical samples ranging from 1 × 10^0^ to 5.2 × 10^6^ parasites per 10^6^ human cells was observed (**Figure 6C**). Depending on the parasite load of the clinical samples and the analytical sensitivity of the newly developed PCR/CRISPR assays, detection of *Leishmania* target DNA was achieved by PCR/CRISPR in different types of clinical specimens that included invasive sample types (biopsies and lesion scrapings using a sterile lancet) and non-invasive sample types (cytology brushes, swabs, and filter paper lesion impressions) (**Supplementary File 2**). Of the 49 clinical specimens analyzed, the majority were biopsies (n = 26; 53.1%), followed by swabs (n = 11; 22.4%). The kDNA qPCR test resulted in 84.6% (22/26) of biopsy specimens, 81.8% (9/11) of swab specimens, 60% (3/5) of cytology brush specimens, 100% (4/4) of scraping specimens, and 33.3% (1/3) of filter paper lesion impressions testing positive for *Leishmania* DNA. The kDNA PCR/CRISPR assay showed 100% concordance with the qPCR test in all specimen types. Of the 39 qPCR positive specimens, *Leishmania* DNA was detected by the 18S PCR/CRISPR assay in 95.5% (21/22) of biopsy specimens, 44.4% (4/9) of swab specimens, 100% (3/3) of cytology brush specimens, and 100% (4/4) of scraping specimens, while no *Leishmania* DNA was detected in the single filter paper lesion impression sample that tested positive by qPCR.

**Figure 6.**
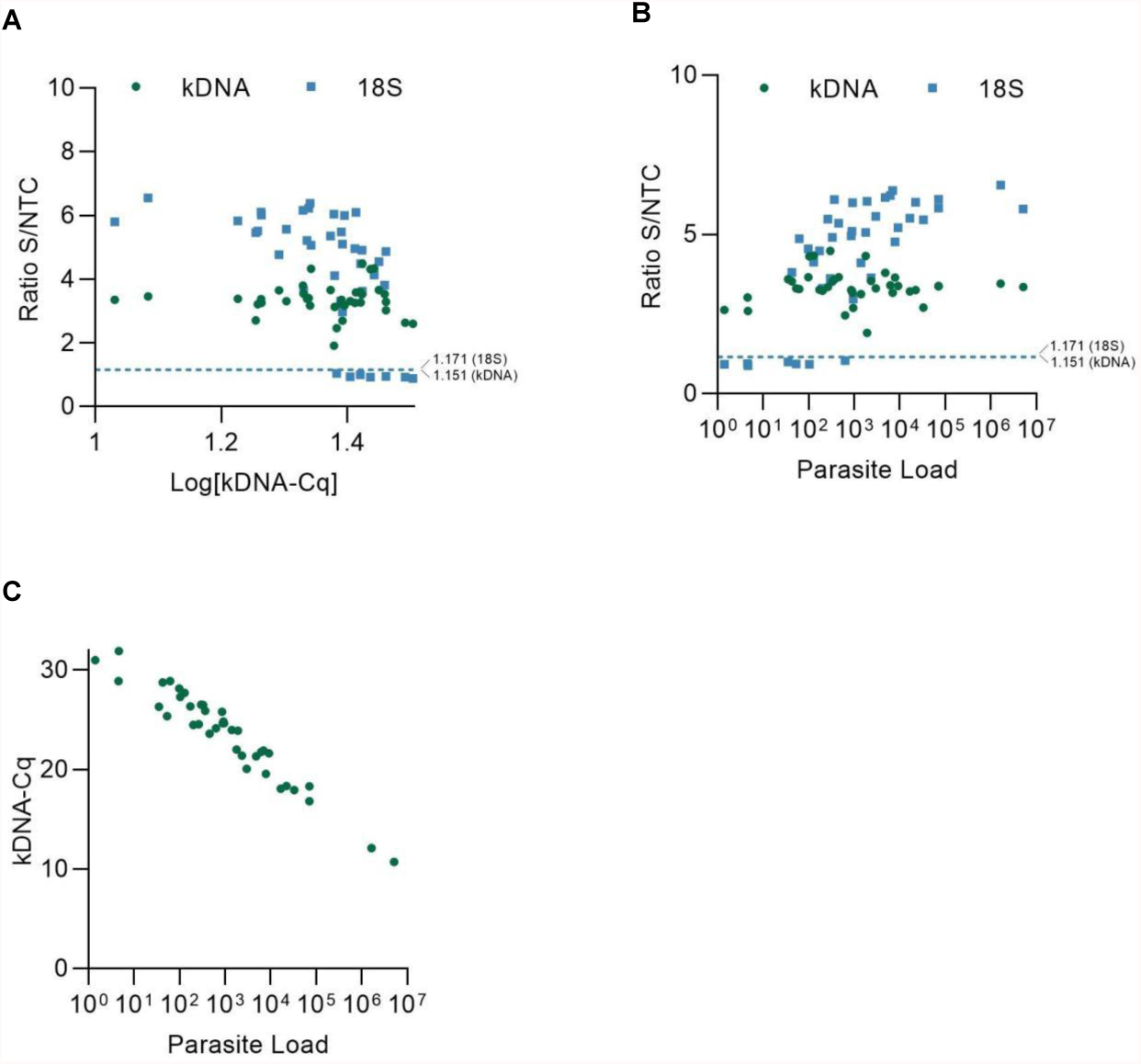
Performance evaluation of PCR/CRISPR assays across the range of parasite load levels in clinical samples. (**A**) Scatter plot showing the fluorescence ratio (fluorescence signal obtained at 20-min time point for kDNA and at 10-min time point for 18S rDNA in the test sample relative to the NTC) from Cas12a reactions for *Leishmania* targets in clinical samples (data shown in Figures 5B,C) *versus* the log of the quantification cycle (Cq) values of the kDNA qPCR assay. (**B**) Scatter plot showing the fluorescence ratio from Cas12a reactions for *Leishmania* targets in clinical samples (data shown in Figures 5B,C) *versus* the parasite load determined by kDNA qPCR. The dashed lines shown in **A**,**B** indicate the positive threshold cutoff value (mean of the fluorescence ratio of negative clinical samples + 3SD; n = 10) for detection of *Leishmania* targets (cutoff value = 1.151 for kDNA and 1.171 for 18S rDNA). (**C**) Scatter plot showing the inverse correlation between Cq values (determined by kDNA qPCR) and parasite load in clinical samples. The parasite load is expressed as the number of *Leishmania* parasites/10^6^ human cells.

As an alternative method to analyze the PCR/CRISPR results on clinical samples, we calculated the percentage positivity (PP) for each sample from the raw fluorescence data and performed a statistical analysis to select the optimal Pr-cutoff/PP to discriminate amongst positive and negative results. We considered the verified PCR/CRISPR results on repeat testing. For the kDNA PCR/CRISPR test, a logistic regression could not be fit due to perfect separation. The predictor variable (kDNA PP) predicted the outcome variable (qPCR result, i.e., measurable or not detectable Cq value) perfectly (**Figure 7A**). Positive samples for *L*. (*Viannia*) kDNA (n = 39) had a PP > 29.9%, whereas negative samples (n = 10) showed a PP ≤ 29.9% (**Figure 7B**). The area under the ROC curve for the kDNA PCR/CRISPR assay was 1 (**Figure 7C**). Regarding the 18S PCR/CRISPR test results, the sets of sensitivity/specificity and their relationship calculated from the logistic regression model using the 18S PP as the predictor variable are shown in **Figure 7D**. Classification of samples based on the Pr-cutoff/PP (18.251%) that maximized the Youden’s J index resulted in 33 samples (67.3%) identified as *Leishmania* positive and 16 samples (32.7%) as negative (**Figure 7E**). One sample (CL-30) had a PP = 18.425% and was classified as positive with the Pr-cutoff/PP, whereas this sample was classified as negative with the cutoff equal to the mean + 3SD. That sample CL-30 had a raw fluorescence value (measured in relative fluorescence units, RFU) of 1118 at the time point 10 min of the Cas12a reaction for the 18S rDNA target, and the fluorescence signal did not show a marked increase over the Cas12a reaction time (RFU at 120 min = 2892), similarly to the negative control with human gDNA (RFU at 120 min = 2764) (**Supplementary File 2**). The 18S PCR/CRISPR assay efficacy, measured by Youden’s J index, was 0.846, and the area under the ROC curve was 0.8795 (**Figure 7F**).

**Figure 7.**
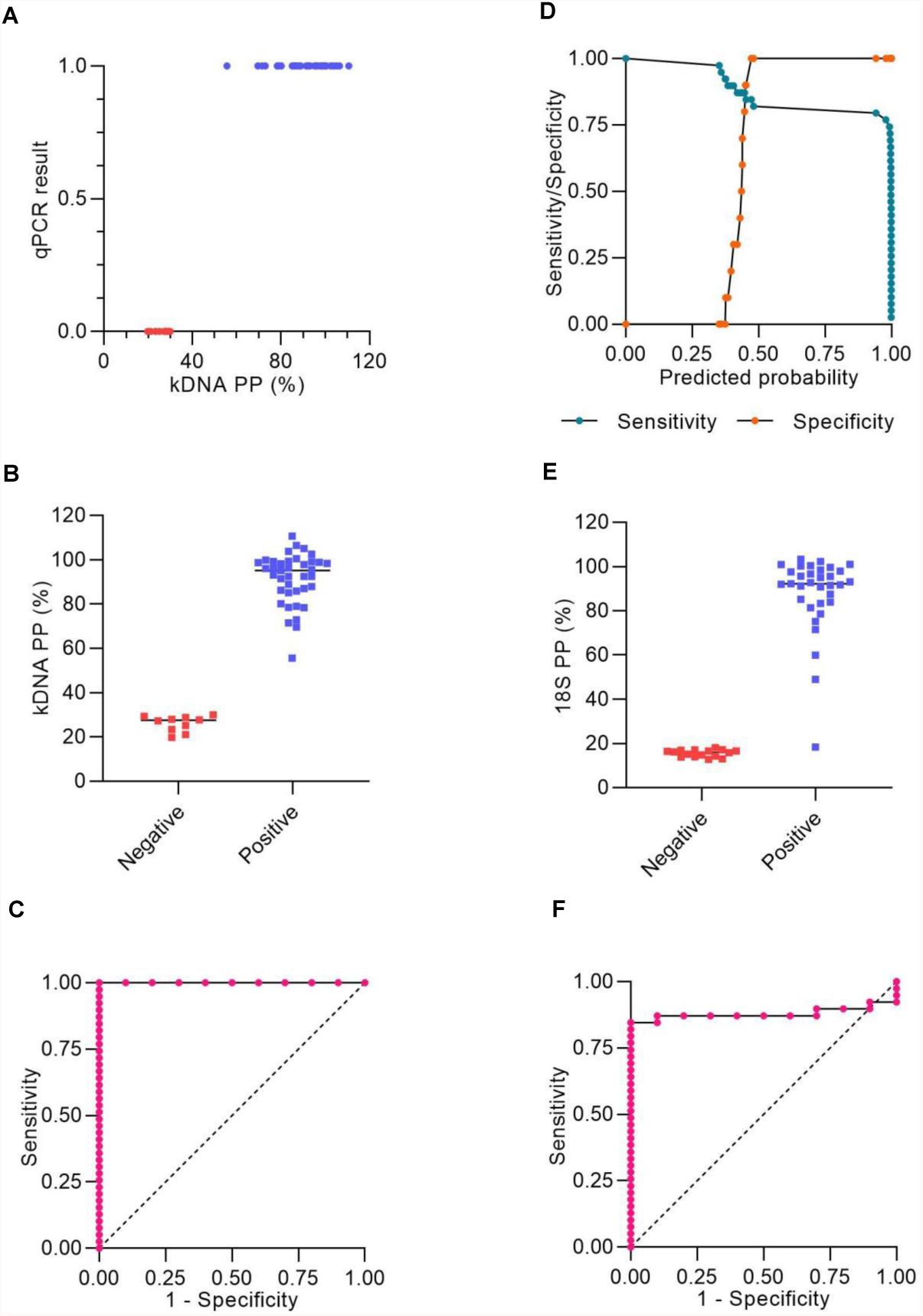
Statistical analysis of Cas12a assay results. (**A-C**) Raw fluorescence data of the Cas12a detection assay on the kDNA target using DNA from clinical samples were expressed as percentage positivity (PP) relative to a positive control, which was run in each plate. (**A**) The predictor variable (PP) predicts the outcome variable Y (qPCR result) perfectly since PP > 29.9% corresponds to Y = 1 (positive) and PP ≤ 29.9% corresponds to Y = 0 (negative). (**B**) Plot representing the PP of positive (n = 39) and negative (n = 10) results obtained by the kDNA PCR/CRISPR assay in clinical samples. (**C**) ROC curve for the kDNA PCR/CRISPR test, ratio of true positive rate (sensitivity) vs. false positive rate (1-specificity) with area under the ROC curve of 1. (**D-F**) Raw fluorescence data of the Cas12a detection assay on the 18S rDNA target gene using DNA from clinical samples were expressed as PP relative to a positive control, which was run in each plate. (**D**) Sensitivity/specificity analysis vs. predicted probability (i.e. probability cutoff). The probability cutoff point determines the sensitivity (85%) and specificity (100%) of the 18S PCR/CRISPR assay. (**E**) Plot representing the PP (cutoff: 18.251%) of positive (n = 33) and negative (n = 16) results obtained by the 18S PCR/CRISPR assay in clinical samples. (**F**) ROC curve for the 18S PCR/CRISPR test, ratio of true positive rate (sensitivity) vs. false positive rate (1-specificity) with area under the ROC curve of 0.8795.

We compared the test performance (considering the verified results on repeat testing) by calculating the positive and negative percent agreement (PPA and NPA, respectively) and Cohen’s kappa relative to the kDNA qPCR comparator method (**Table 1**). The kDNA PCR/CRISPR test correctly identified all 39 samples that had tested positive using the kDNA qPCR test, with 100% PPA (95% CI = 91.0% to 100.0%). Cohen’s kappa was 1, consistent with perfect agreement between methods. Using the classification cutoff equal to the mean + 3SD, the 18S PCR/CRISPR test had a PPA of 82.1% (95% CI = 67.3% to 91.0%) and Cohen’s kappa was 0.65, which represents a substantial strength of agreement. Based on the Pr-cutoff/PP, the PPA for the 18S PCR/CRISPR test was 84.6% (95% CI = 70.3% to 92.8%) and Cohen’s kappa was 0.69. The PPA for the kDNA cPCR test was 92.3% (95% CI = 79.7% to 97.3%) and Cohen’s kappa was 0.83, consistent with excellent agreement between the kDNA cPCR and qPCR methods. Keeping in mind the limitation of having analyzed a relatively small sample set, particularly samples that were negative for *Leishmania* infection with the reference test (n = 10), the NPA of all evaluated tests was 100% (95% CI = 72.2% to 100.0%).

**Table 1.** Concordance analysis of molecular tests compared to the reference kDNA qPCR test for detection of *Leishmania* in clinical samples. The PPA/NPA agreement measures between the reference method (kDNA qPCR) and any of the other tests, and Wilson score 95% confidence intervals (CI) were calculated using the Analyse-it software for Microsoft Excel. The Kappa coefficient and 95% CIs were calculated as described previously by McHugh (2012) (https://www.ncbi.nlm.nih.gov/pmc/articles/PMC3900052/). PPA, positive percent agreement. NPA, negative percent agreement. SD, standard deviation. Pr-cutoff, probability cutoff. PP, percentage positivity. cPCR, conventional PCR. qPCR, real-time quantitative PCR.

| Classification threshold / readout | Test | kDNA qPCR |  | PPA | NPA | Kappa |
| --- | --- | --- | --- | --- | --- | --- |
|  |  | positive | negative | % (95% CI) | % (95% CI) | (95% CI) |
| specific detectable band (gel electrophoresis) | kDNA cPCR |  |  |  |  |  |
|  | positive | 36 | 0 | 92.3 (79.7-97.3) | 100.0 (72.2-100.0) | 0.83 (0.55-1.11) |
|  | negative | 3 | 10 |  |  |  |
| mean + 3SD | kDNA PCR/CRISPR |  |  |  |  |  |
|  | positive | 39 | 0 | 100.0 (91.0-100.0) | 100.0 (72.2-100.0) | 1.00 (0.72-1.28) |
|  | negative | 0 | 10 |  |  |  |
| mean + 3SD | 18S PCR/CRISPR |  |  |  |  |  |
|  | positive | 32 | 0 | 82.1 (67.3-91.0) | 100.0 (72.2-100.0) | 0.65 (0.39-0.91) |
|  | negative | 7 | 10 |  |  |  |
| Pr-cutoff/PP | positive | 33 | 0 | 84.6 (70.3-92.8) | 100.0 (72.2-100.0) | 0.69 (0.42-0.96) |
|  | negative | 6 | 10 |  |  |  |
|  | Total | 39 | 10 |  |  |  |

Lastly, we carried out an evaluation of the intermediate precision (within-laboratory reproducibility or inter-assay variability) of measurements across PCR amplification and Cas12a-based detection runs for the positive and negative controls included alongside test samples. While the measurements were performed by a single operator using the same instrument (Synergy H1 plate reader), variable conditions across runs were the time scale (5 months) and reagent batches (i.e., two batches of *in vitro* transcribed crRNAs). For both the kDNA and 18S PCR/CRISPR assays, the time course data of Cas12a reactions showed a steady increase in fluorescence for the positive control (PC), whereas the fluorescence signal remained low in the HC and NTC controls of PCR and CRISPR reactions (**Figures 8A** and **8D**), as expected. For the kDNA target, the analysis of the PC showed increased variability (standard deviation) of mean raw fluorescence values over the reaction time course across runs (n = 12) (**Figure 8A**). Normalization of the fluorescence signal relative to the NTC (i.e., fluorescence ratio) showed a similar trend in the overall distribution (spread) of data values (**Figure 8B**). At the selected 20-min time point of the Cas12a reaction on the kDNA target for data analysis, there was a clear distinction between specific and background signal: the PC showed fluorescence ratio values ranging from 2.08 to 5.71 and a median of 3.48, while the negative controls yielded a maximal fluorescence ratio of 1.28 (**Figure 8C**). For the 18S rDNA target, raw fluorescence measurements for the PC showed a mostly similar variability over the reaction time course across runs (n = 9) (**Figure 8D**). The determined fluorescence ratio values for the PC showed less variability at early time points (e.g. at 10 and 20 min) as compared to later time points (40 and 60 min) of the Cas12a reaction employing the 18S-crRNA (**Figure 8E**). At the selected 10-min time point of the Cas12a reaction on the 18S rDNA target for data analysis, the fluorescence ratio clearly distinguished specific from background signal: the PC showed fluorescence ratio values ranging from 5.00 to 6.92 and a median of 5.67, while the negative controls showed a maximal fluorescence ratio of 1.32 (**Figure 8F**). Altogether, these data showed that the positive and negative controls provided consistent reference data points in each experiment.

**Figure 8.**
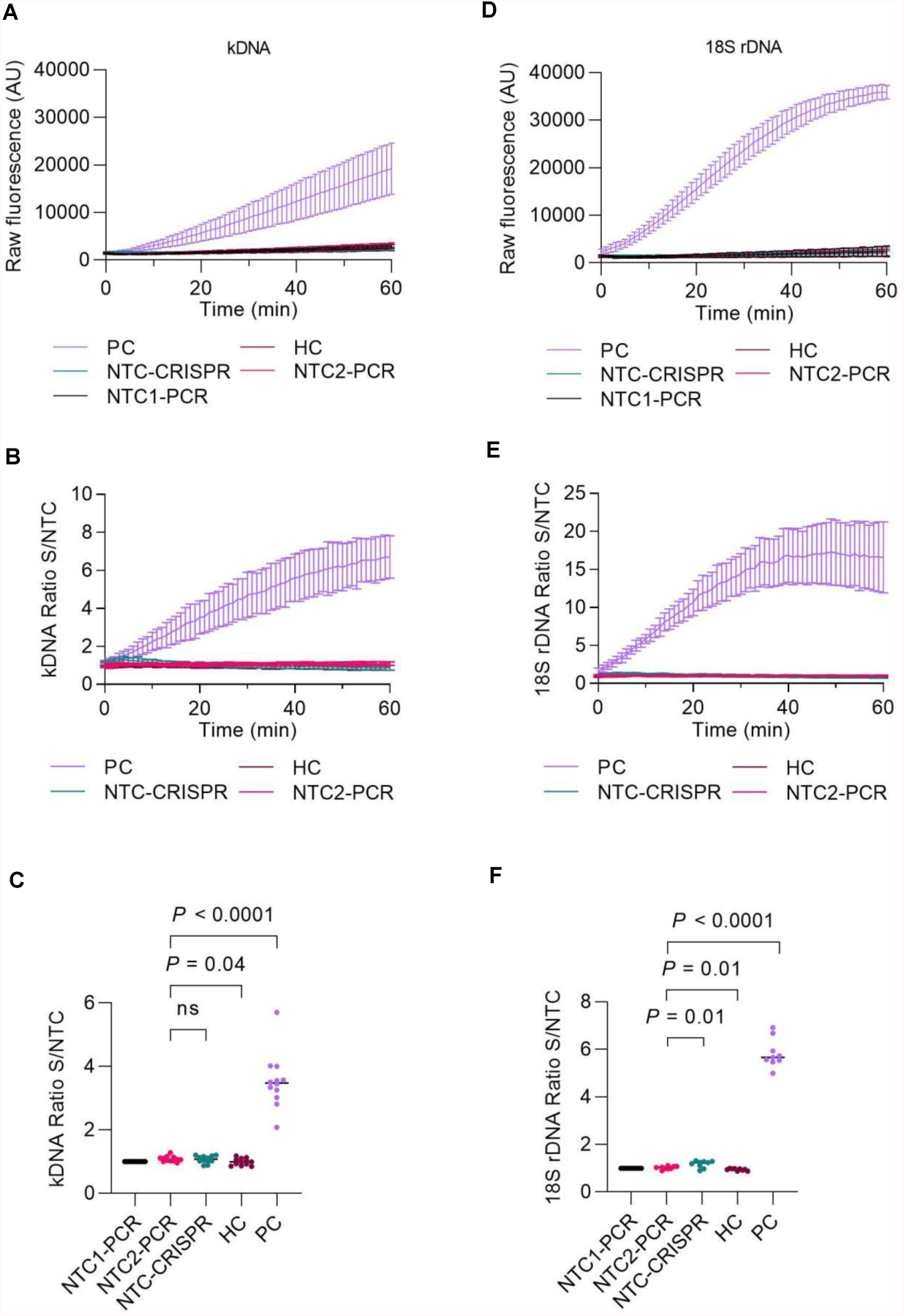
Assessment of PCR/CRISPR assay precision for detection of *Leishmania* targets. Measurements in independent experiments of the positive and negative controls (included alongside test samples) allowed to assess the intermediate precision (within-laboratory reproducibility) across PCR amplification and Cas12a-based detection runs. Fluorescence measurements taken on the Synergy H1 plate reader were considered for this analysis. The assays were performed by the same operator over a period of 5 months; two reagent batches of *in vitro* transcribed crRNAs were used. The positive control (PC) consisted of gDNA from *L*. (*V*.) *braziliensis* M2904 (5 × 10^4^ parasite genome equivalents/reaction used as input DNA). The human negative control (HC) consisted of gDNA from PBMC of a healthy donor (40 ng of input DNA). Two NTC reactions were included in PCR runs to monitor for contamination: NTC1, kept closed without water addition [this control served for normalization of raw fluorescence values of the Cas12a reaction, expressed as a fluorescence ratio], and NTC2, made with water instead of the template. NTC-CRISPR, no-template control of the Cas12a reaction. (**A**) Raw fluorescence signal from the Cas12a reaction over 1 h for the kDNA target. Mean raw fluorescence values (arbitrary units, AU) ± SD are plotted as lines with error bars (n = 12). (**B**) The data in **A** were normalized to the fluorescence of the NTC1 (PCR blank) and expressed as a fluorescence ratio. (**C**) The normalized data in **B** were used for detection analysis at the 20-min time point. (**D**) Raw fluorescence signal from the Cas12a reaction over 1 h for the 18S rDNA target. Mean raw fluorescence values (arbitrary units, AU) ± SD are plotted as lines with error bars (n = 9). (**E**) The data in **D** were normalized to the fluorescence of the NTC1 (PCR blank) and expressed as a fluorescence ratio. (**F**) The normalized data in **E** were used for detection analysis at the 10-min time point. For panels **C, F**, pairwise comparisons to the NTC2 (PCR blank) were done using an unpaired *t*-test. Two-tailed *P* values are shown; non-significant *P* values (*P* > 0.05) are indicated by ns.

## DISCUSSION

Accurate detection of *Leishmania* infections is critically important in the confirmatory diagnosis of leishmaniasis. In the Americas, where tegumentary leishmaniasis is highly endemic, skin lesions suspicious of CL may be caused by other infectious agents like mycobacteria or fungi that are prevalent in the same endemic areas or non-infectious conditions like skin neoplasms (Tirelli et al., 2017). Differential diagnosis of TL requires reliable confirmatory tests that help to guide clinical management and appropriate treatment, to avoid exposing patients to unnecessary and toxic antileishmanial drugs (Burza et al., 2018). In this study, we report the development of two novel CRISPR-Cas12a-based assays for the detection of *Leishmania* spp. in human clinical specimens. We present the analytical validation using laboratory reference strains and the assessment of the performance of the assays in a panel of clinical samples with PCR-predetermined *Leishmania* infection status derived from patients with suspected CL from Cusco, an endemic region where *L*. (*Viannia*) parasites circulate, with predominance of *L*. (*V*.) *braziliensis* and less frequently *L*. (*V*.) *guyanensis* and *L*. (*V*.) *lainsoni* infections (Lucas et al., 1998; Sandoval-Juárez et al., 2020). The choice of the molecular target sequences was based on their multicopy nature, conservation across species of *Leishmania* at the genus level (18S rDNA) to allow pan-*Leishmania* detection or at the *L*. (*Viannia*) subgenus level (kDNA minicircle conserved region) to detect the main parasite species that cause TL in the Americas, and the fact that they are well-validated genetic targets widely used for the molecular diagnosis of leishmaniasis (Van der Auwera and Dujardin, 2015; Akhoundi et al., 2017).

The 18S rDNA locus is a region on chromosomal DNA that is highly conserved across *Leishmania* species (van Eys et al., 1992), present at about 10–170 copies per genome (Leon et al., 1978; Inga et al., 1998), that has been harnessed for the development of *Leishmania* detection assays of high sensitivity (de Paiva Cavalcanti et al., 2013; Adams et al., 2014; León et al., 2017; Filgueira et al., 2020; Rosales-Chilama et al., 2020) as well as for the estimation of parasite loads (Bezerra-Vasconcelos et al., 2011; Filgueira et al., 2020). Here, the estimated analytical sensitivity of our 18S PCR/CRISPR assay with DNA of *L. braziliensis* M2904 culture promastigotes corresponded well with its performance in clinical samples being capable of detecting the equivalent of 1 parasite per reaction. This detection capability of our novel assay is similar to a previously reported 18S qPCR assay (Rosales-Chilama et al., 2020), which was able to detect down to 10^−1^ *L*. (*V*.) *panamensis* promastigotes and 1 intracellular amastigote per reaction. Our assay enabled detection of species of both *L*. (*Leishmania*) and *L*. (*Viannia*) subgenera, with no cross-reaction with *T. cruzi* DNA (Y strain) or human DNA. Due to the high sequence conservation of 18S rDNA throughout the Trypanosomatidae, other non-*Leishmania* trypanosomatids not evaluated here could probably be detected with the 18S PCR/CRISPR assay. A low specificity of the 18S rDNA target was indeed the case with a 18S qPCR assay that was not exclusive for *Leishmania* detection because it showed cross-reactivity with non-*Leishmania* trypanosomatids, including *T. cruzi* (Y strain), *T. rangeli* and lower insect-dwelling trypanomatids (Filgueira et al., 2020).

The minicircles of kinetoplast (mitochondrial) DNA constitute the most widely used genetic target for extremely sensitive *Leishmania* detection (Akhoundi et al., 2017), because they are present in high copy numbers, at about 10,000 copies per cell (Simpson, 1987). qPCR assays that target kDNA minicircles proved highly sensitive and accurate for detection and quantification of *Leishmania* spp. in human clinical specimens (Weirather et al., 2011; Jara et al., 2013). Here, we tested two kDNA primer sets for the preamplification step, because differences in the performance of kDNA primer sets for detection of *Leishmania* spp. have been reported (Weirather et al., 2011) and likely reflect the heterogeneity in minicircle sequence classes between species and even between strains of the same species (Simpson, 1997; Ceccarelli et al., 2014; Kocher et al., 2017). The analytical sensitivity on DNA from *L. braziliensis* promastigotes with either kDNA primer set in combination with the examined kDNA-crRNA reached down to 5 × 10^−2^ parasite equivalents per reaction, which approximated that of a previously validated kDNA qPCR assay for *L*. (*Viannia*) detection (5 × 10^−3^ parasite equivalents per reaction; Jara et al., 2013). The combination of the kDNA primer set 1 (PCR preamplification) and the kDNA-crRNA (Cas12a assay) was found to be specific for *L*. (*Viannia*) detection, and thus it was further tested with clinical samples. The kDNA PCR/CRISPR assay did not show cross-reactivity with *T. cruzi* Y strain or human DNA.

The CRISPR-Cas12a-based assays developed here showed good performance in detecting *Leishmania* DNA targets across a wide range of parasite loads in clinical specimens that included both invasive and non-invasive sample types. Remarkably, the kDNA PCR/CRISPR assay showed perfect agreement with the reference kDNA qPCR assay in classifying the samples with positive (detected) or negative (not detected) *Leishmania* infection status. Of the 39 samples being positive by the kDNA PCR/CRISPR and qPCR assays, 7 samples were negative by the 18S PCR/CRISPR assay, a difference related to the lower copy number of 18S rDNA. These 7 samples contained the equivalent of less than one parasite per reaction corresponding to the low range of parasite loads (**Figure 6B** and **Supplementary File 2**). An important aspect to stress is that because of the high copy number of kDNA minicircles and therefore the high detection sensitivity achieved with this target, the kDNA PCR/CRISPR assay was more prone to false-positive results. If we would not have carried out assay repeat testing to validate the results, the false positive rate of the kDNA PCR/CRISPR assay was 4.1% (i.e. 2 false positives of 49 patient samples), showing 95.9% accuracy (47/49 test results in agreement with kDNA qPCR results). This drawback with the kDNA target was observed in another study using minicircle kDNA qPCR, in which false-positive reactions were reduced by combining the kDNA target with 18S rDNA in a multiplex probe-based qPCR assay and setting a cutoff Cq value to exclude false-positive samples (Eberhard et al., 2018). Similar to our study, the possibility of carryover contamination is a challenge with any of the two-step CRISPR-based detection platforms (Kellner et al., 2019; Nguyen et al., 2022). All these comprise a first step of target preamplification, either by PCR [as in HOLMES (Li et al., 2018)] or isothermal amplification methods [as in SHERLOCK (Gootenberg et al., 2017), DETECTR (Chen et al., 2018), and ENHANCE (Nguyen et al., 2020; Nguyen et al., 2022)], followed by the CRISPR-based detection reaction. This implies multiple manual operations and opening the lid of the amplification reaction tubes to transfer a volume of preamplified target DNA for the detection phase of the assay, thereby increasing the risk of amplicon carryover contamination. To prevent this, proper laboratory practices and reaction setup must be in place, with physically separated laboratory work areas in which pre- and post-amplification manipulations are performed. As followed here, we recommend processing patient samples in groups of 10-12 per run alongside appropriate negative and positive controls of the amplification and detection reactions to ensure reliability and validity of the assays. The human RNase P gene was used as sample control to verify an efficient DNA extraction and validate true-negative results (i.e. those samples with undetectable *Leishmania* DNA but positive for human RNase P). We also recommend to perform two replicate assays to validate the test results in clinical samples as well as performing rigorous data quality control as presented here. Combining preamplification and CRISPR-based detection in a single one-pot assay such as with one-pot SHERLOCK (Gootenberg et al., 2018; Kellner et al., 2019), AIOD-CRISPR (Ding et al., 2020), and other innovative ways (Yin et al., 2020; Wang et al., 2021; Hu et al., 2022) has been shown to be faster, simpler, amenable to large-scale and quantitative applications, and less prone to contamination, albeit with lower sensitivity and a more challenging assay optimization phase than with two-step CRISPR-based assay formats. In fact, a recent study that exploited the Cas13a-based SHERLOCK method to develop an assay for detection of the malaria-causing *Plasmodium* parasites (Cunningham et al., 2021) was unable to replicate the one-pot reaction format previously described for viral targets (Gootenberg et al., 2018), achieving instead the best performance with a two-step reaction format. Another way to greatly reduce carryover contamination is the use of dUTP instead of dTTP in the amplification reaction master mix and treating subsequent reactions with uracil DNA glycosylase (UDG) prior to PCR or isothermal amplification. UDG will degrade dUTP-containing amplicons, thus preventing these molecules to serve as templates (Longo et al., 1990). Integration of LAMP amplification with UDG digestion followed by downstream CRISPR-based detection was successful at lowering the risk of false positives (Qian et al., 2019; Nguyen et al., 2022), though this can result in a higher limit of detection of the method (as found with ENHANCEv2, Nguyen et al., 2022).

This study has a main limitation. This concerns the sample size of tested clinical samples, particularly the small number of samples whose *Leishmania* infection status was negative with the reference qPCR test (n = 10). However, this study was designed to provide proof-of-concept of the newly developed CRISPR-based assays for application with patient samples. Keeping in mind the limitation of sample size, we did not calculate estimates of sensitivity and specificity (diagnostic accuracy) and used instead inter-rater agreement measures.

We used standard PCR as the target preamplification strategy to provide proof-of-concept CRISPR-based assays for highly sensitive and specific detection of *Leishmania* infection in clinical specimens. Our assays utilize a detection method similar to HOLMES (a one-hour low-cost multipurpose highly efficient system), which uses LbCas12a-mediated DNA detection combined with PCR preamplification to achieve high sensitivity (Li et al., 2018). The main differences between our methods concern the PCR amplification cycles (35 cycles for ∼45 minutes in HOLMES vs. 45 cycles for ∼90 minutes in our study) and the incubation temperature of the Cas12a assay (37 °C in HOLMES vs. 25 °C in our study). Our preliminary assays indicated that shortening the period of PCR preamplification to 35 cycles also enriched the target molecules to enable sufficiently high detection sensitivity, though more evaluations are needed to assess that parameter in the assay performance using clinical samples. While the LbCas12a enzyme displays optimal *trans*-cleavage activity at 37 °C (Chen et al., 2018), it is also functional at room temperature (25 °C) (Ding et al., 2020; Xiong et al., 2020), though with a slower enzyme kinetics at 25 °C in a one-pot assay format (Ding et al., 2020) compared to a two-step detection workflow (Xiong et al., 2020). Here, we performed the LbCas12a-based detection assays at 25 °C to facilitate further development and adaptation of the assays for POC applicability and observed a good performance (high signal-to-noise ratios).

Nowadays, the PCR technique is widely used in research and reference laboratories in Latin America for molecular diagnosis of TL (Moreira et al., 2018). Accordingly, our assays in their current format have the potential for widespread application in laboratory settings and reference centers that have the needed infrastructure and resources to perform PCR and analyze Cas12a assay results with a fluorescence plate reader. Further studies are needed to extend the assessment of the analytical and clinical performance of both CRISPR-based assays to different endemic regions across the Americas where different *Leishmania* species and parasite variants circulate. We envisage that our 18S PCR/CRISPR assay would be useful for detection of all medically relevant *Leishmania* species in clinical specimens containing the equivalent of at least 1 parasite/reaction, such as in cutaneous lesions of recent onset (≤ 3 months) that harbor higher parasite loads than lesions with longer evolution (Weigle et al., 2002; Jara et al., 2013). Furthermore, this novel 18S CRISPR-based assay could be of global applicability if validated on a broader scale in different geographic regions where different forms of leishmaniasis are endemic. The kDNA PCR/CRISPR assay would be a useful addition to specifically detect New World *L*. (*Viannia*) infections, including those at low parasite load levels, as in chronic skin lesions and in mucosal lesions (Jara et al., 2013), which are undetectable by microscopy (Weigle et al., 2002). Both newly developed CRISPR-based assays provide positive or negative results for *Leishmania* detection, but do not enable quantitative applications such as parasite load quantification, as can be determined by real-time qPCR assays (Moreira et al., 2018). Altogether, the results shown here support the potential applicability of the novel 18S and kDNA PCR/CRISPR assays for first-line diagnostic purposes that require detection of *Leishmania* infections at the genus and *L*. (*Viannia*) subgenus levels, respectively.

Several molecular methods that employ different genetic targets have been developed and are in use for molecular detection of *Leishmania*, usually at the genus or subgenus level (Akhoundi et al., 2017). PCR-based methods, particularly real-time qPCR assays, have proved of clinical utility for the detection of *Leishmania* infections and quantification of parasite load in host tissues (Moreira et al., 2018). Many studies report in-house assays using varied protocols involving different types of specimens, which makes it difficult to compare results between different studies. The need for standardization and proper validation of a real-time PCR-based methodology for the molecular diagnosis of TL in the Americas has been underscored (Moreira et al., 2018; Rosales-Chilama et al., 2020; Filgueira et al., 2020) and efforts toward this respect are ongoing (Moreira et al., 2018; Filgueira et al., 2020). This is expected to provide a consensus well-established methodology for broad use at research and reference laboratories across Latin America. There is also the critical need for accessible, sensitive and accurate diagnostic tools that offer easy operation and shorter turnaround time for results to diagnose TL in low-resource settings in endemic regions. CRISPR-based diagnostics has the potential to contribute to this and to become important tools for parasite detection in the clinical and epidemiological context, as illustrated in two recent studies that developed SHERLOCK CRISPR-based assays applied to *Plasmodium* detection and species identification (Lee et al., 2020; Cunningham et al., 2021) as well as drug-resistance genotyping (Cunningham et al., 2021). One advantage of CRISPR-based detection methods over other molecular techniques used in infectious disease diagnosis is the greater flexibility and versatility of the former with respect to simplifying and optimizing the assay workflow as well as adapting it to user-friendly readouts for POC use (at/near the site of patient care), making these tools accessible at the primary health care level in resource-limited settings. In this scenario, our CRISPR-based assays are amenable to further optimization and adaptation for use in POC settings. We foresee different options. The assays can be transferred to low-cost and portable PCR (e.g. Bento Lab^®^, Alcántara et al., 2021a) and fluorescence reader (Katzmeier et al., 2019; Lee et al., 2020; Xiong et al., 2020) devices. A simplified workflow for *Leishmania* detection can be optimized starting from the sample preparation step (e.g. evaluating rapid means to extract nucleic acids with chelating agents to inactivate nucleases), followed by introducing an isothermal amplification approach coupled to downstream Cas12a-based detection with lateral flow strip readout vs. the use of a handheld fluorimeter, as exemplified by a recent study on field-applicable *Plasmodium* detection (Lee et al., 2020). Experimental strategies where *Leishmania* amastigote cells can firstly be enriched from a patient’s lesional tissue sample using affinity reagents for cell capture (Chen, 2019; Gray et al., 2020) during the process of sample preparation are worth to be explored for a diagnostic POC test, given the usually low parasite load found in CL lesions. Among the isothermal NAATs, LAMP and recombinase polymerase amplification (RPA) are commonly used in the literature, since both exhibit features that are ideal for POC settings. We find LAMP better suited for field use in TL endemic regions across Latin America, as LAMP reagents are easier and cheaper to get (the RPA commercial kit is supplied by the single manufacturer, TwistDx™). Moreover, LAMP is more sensitive than RPA (Parida et al., 2006) and can be integrated into a low-tech portable heating device (Curtis et al., 2012). On the other hand, LAMP primer design is more complex because 4-6 primers are required in a single LAMP reaction to facilitate amplification at multiple sites of the target DNA. The use of several primers and high concentration of the reaction components (primers, Mg^2+^, dNTPs, and Bst DNA polymerase) makes LAMP more prone to non-specific amplification (Wang et al., 2015). Coupling LAMP or RPA to downstream crRNA-based target detection improves specificity (Kaminski et al., 2021). Both isothermal reactions as well as CRISPR reactions are compatible with lyophilization (freeze drying). Hence, the impact of lyophilizing the reaction reagents on the performance of our assays needs also to be assessed in future work, since lyophilized reagents have improved stability for transport and long-term storage at ambient temperature, reduce the handling steps and contamination risk, thus facilitating nucleic acid POC testing (Lee et al., 2020; Nguyen et al., 2022). Furthermore, the lyophilized reagents and assay controls can be integrated on a test kit format. Recent advances in CRISPR-based nucleic acid detection toward POC application have focused on engineering CRISPR-Cas systems and optimizing assay conditions to tune the kinetic properties of Cas enzymes, resulting in enhanced detection sensitivity of targets without the need for preamplification and enabling quantitative diagnostic applications (Nguyen et al., 2020; Fozouni et al., 2021; Nalefski et al., 2021). In the future, field testing and validation of the streamlined CRISPR-based assay workflow in the chosen POC test format will be an important step before the test can be implemented and used clinically.

Concerning the development of CRISPR-based biosensors applied to *Leishmania* DNA detection, in a recently published study, Bengtson *et al*. describe a newly developed CRISPR-dCas9-based DNA detection scheme targeting *Leishmania* kDNA minicircles and its potential for instrument-free POC diagnosis of VL in endemic, resource-limited settings (Bengtson et al., 2022). This DNA detection system operates entirely at room temperature (23 °C) and combines isothermal target amplification by RPA and dCas9-based target DNA recognition with a subsequently primed rolling circle amplification (RCA) reaction. The RCA products were designed to fold into G-quadruplex (G4) structures that can associate with hemin to form a peroxidase-mimicking G4 DNAzyme that produces the final colorimetric readout that is visible to the naked eye. This method was combined with instrument-free simplified DNA extraction procedures for blood and urine samples and it demonstrated to work well for highly sensitive detection of target DNA in simulated patient samples. Notwithstanding that this DNA detection scheme was designed to be directly suited for field use, its performance in clinical samples is not yet determined. Interestingly, Bengtson *et al*. chose a conserved 23-mer target sequence within kDNA minicircles that has a PAM site for dCas9 binding, which overlaps with our selected Cas12a crRNA target site (**Figure 2B**). Our chosen target sequence is highly conserved among New World *L*. (*Viannia*) species, whereas the target sequence selected by Bengtson *et al*. is conserved among some members of the *L*. (*Leishmania*) subgenus, namely *L*. (*L*.) *donovani, L*. (*L*.) *infantum* (both etiologic agents of VL in the Old World), *L*. (*L*.) *infantum* (syn. *L. chagasi*) (etiologic agent of VL in the New World), and *L*. (*L*.) *major* (etiologic agent of CL in the Old World), based on *in silico* multiple sequence alignments. While our studies use different CRISPR-based detection approaches, both underscore the great potential of these methods to be adopted for leishmaniasis diagnosis, surveillance and research applications. The performance of the CRISPR-based detection assays/scheme shown in both proof-of-principle studies lays the groundwork for future studies to further develop, adapt and validate these molecular detection systems in clinical and research settings in which the tests are to be used.

## AUTHOR CONTRIBUTIONS

ED, PM, and VA designed the study. JAN and LC-S performed the bioinformatics analyses. MC recruited patients and carried out sampling of lesions. ED and PH performed the experiments. ED, VA, JAN, LC-S, PH, JA, and PM analyzed the data. ED and VA wrote the manuscript with input from all authors. All authors read and approved the final manuscript.

## ACKNOWLEDGMENTS

We thank Roberto Alcántara and Katherin Peñaranda for sharing the CRISPR-Cas12a-based detection assay protocol; Gabriel Mendoza for sharing recombinant LbCas12a; Manuela Verástegui for providing genomic DNA from the *Trypanosoma cruzi* Y strain; and Mirko Zimic for advice on the statistical methods to determine the optimal cutoff value (decision threshold) of a diagnostic test.

