## Supplementary Figures for "Novel CRISPR-based detection of *Leishmania* species"

for

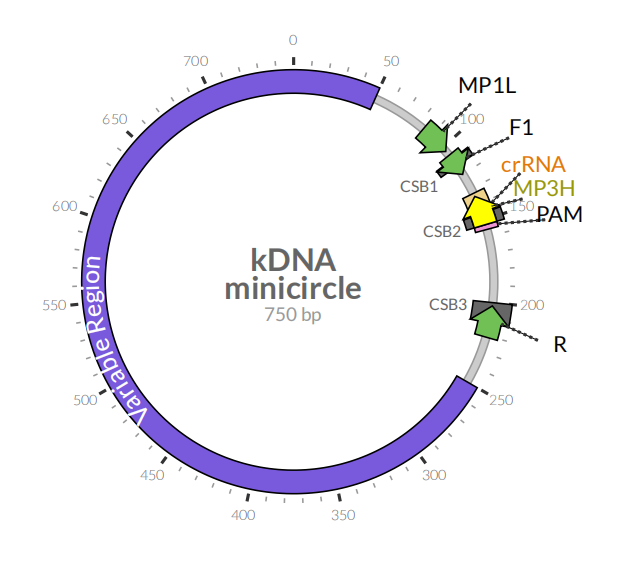
**Figure S1.** Structural organization of *Leishmania* kDNA minicircles (based on the *L.* (*V*.) *braziliensis* M2904 strain, GenBank accession number KY698821). The three conserved sequence blocks (CSB1-3) (Kocher et al., 2017; Camacho et al., 2019) are indicated in grey boxes. Green arrows indicate the binding sites of primers used to amplify a 116 bp-long fragment (primer set 1: F1 and R) or a 138 bp-long fragment (primer set 2: MP1L and R) within the conserved region (those primer pairs were tested in the PCR-based preamplification step prior to Cas12a-based detection). The location of the crRNA guide sequence, which is complementary to the target DNA, and the adjacent PAM sequence in the target DNA are indicated. The primer pair MP1L and MP3H has been previously validated for kDNA cPCR (López et al., 1993) and qPCR (Jara et al., 2013). The AngularPlasmid (<http://angularplasmid.vixis.com/index.php>) was used to prepare this figure.

(C)

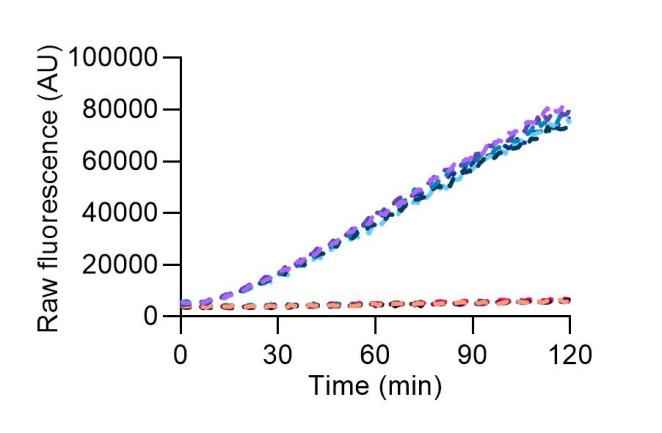

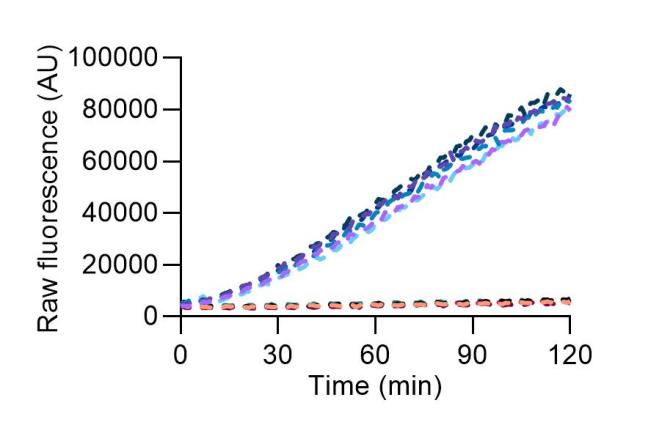

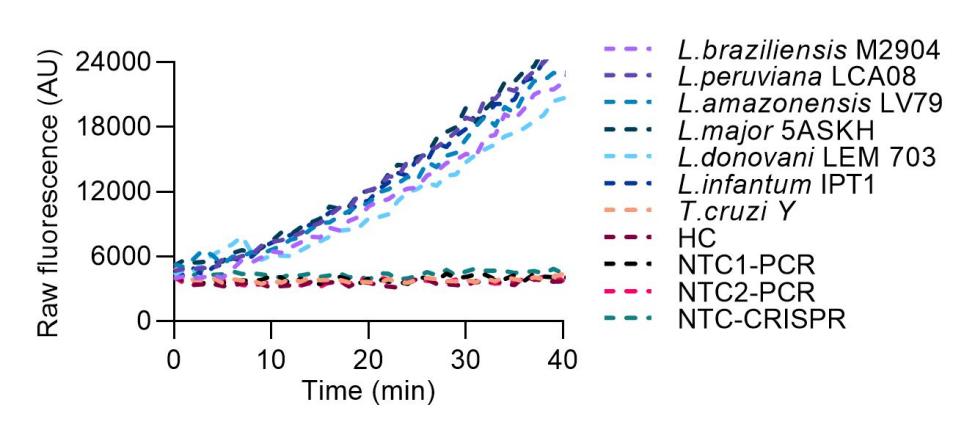

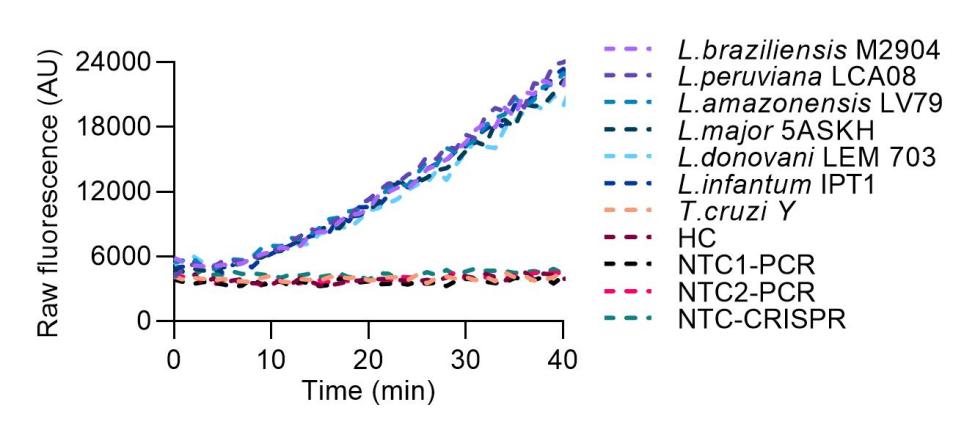

**A**

**B**

**C**

**D**

**Figure S2.** The kDNA primer set 2 was tested in PCR reactions followed by crRNA-guided Cas12a-based detection for specificity assessment of the kDNA PCR/CRISPR assay using DNA from 2 *L*. (*Viannia*) and 4 *L*. (*Leishmania*) strains representative of different *Leishmania* species, and the *T. cruzi* Y strain. Duplicate PCR reactions were amplified in parallel (**A**: replicate 1, **C**: replicate 2) followed by Cas12a detection. Raw fluorescence signal (arbitrary units, AU) over 2 h from Cas12a reactions is shown. (**B**) Same dataset as in **A**, but zoomed to the first 40 min of the Cas12a assay. (**D**) Same dataset as in **C**, but zoomed to the first 40 min of the Cas12a assay. Negative controls included human PBMC gDNA and NTC controls of PCR and CRISPR reactions. Fluorescence measurements in this figure were taken on the Cytation 5 plate reader.

**G**

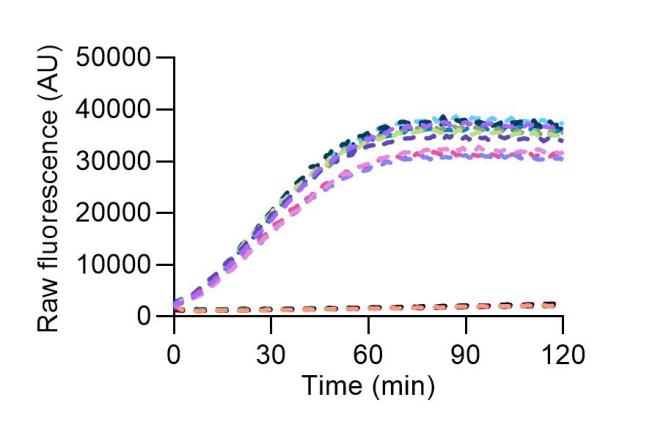

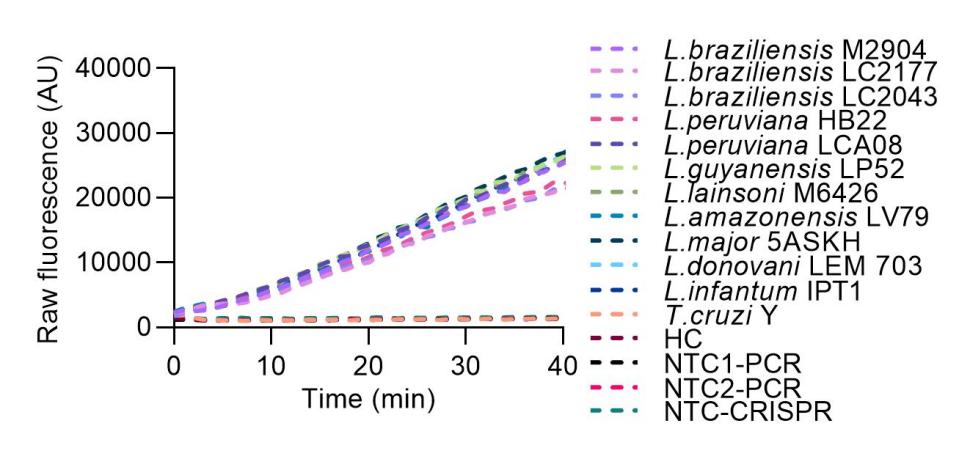

**H**

kDNA - primer set 1

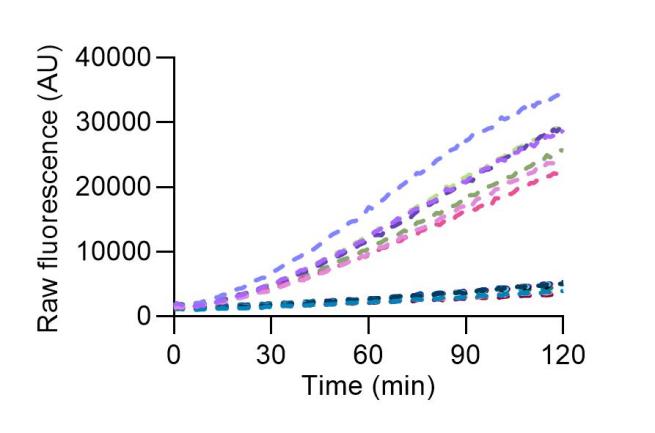

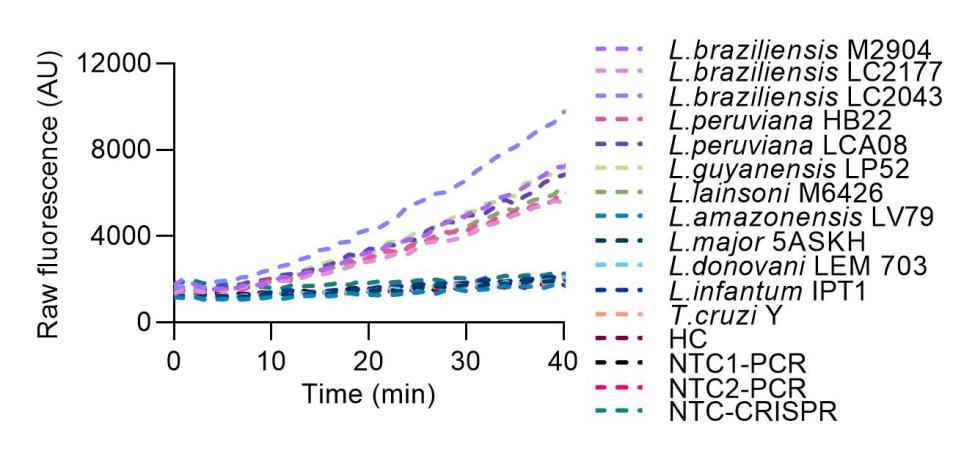

**A**

**B**

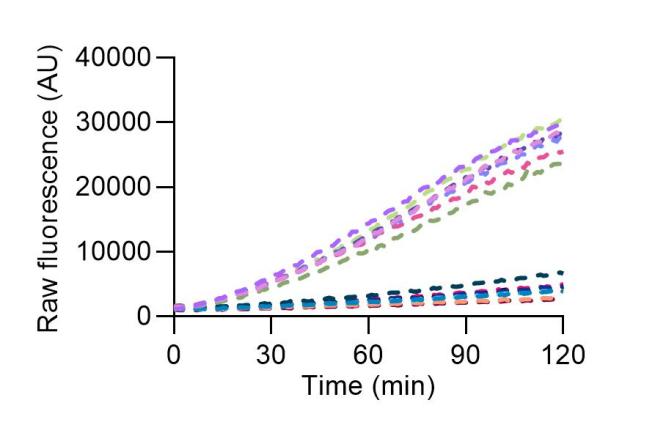

**C**

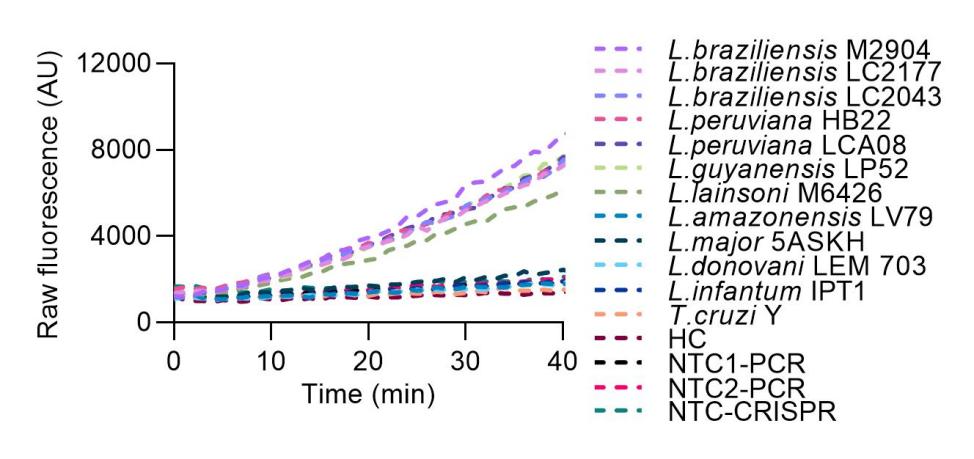

**D**

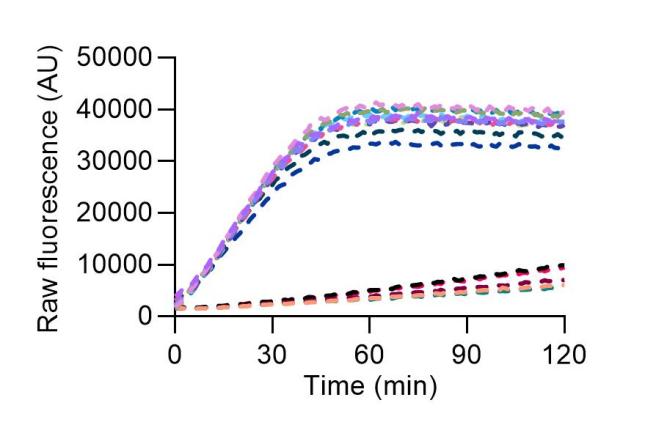

**E**

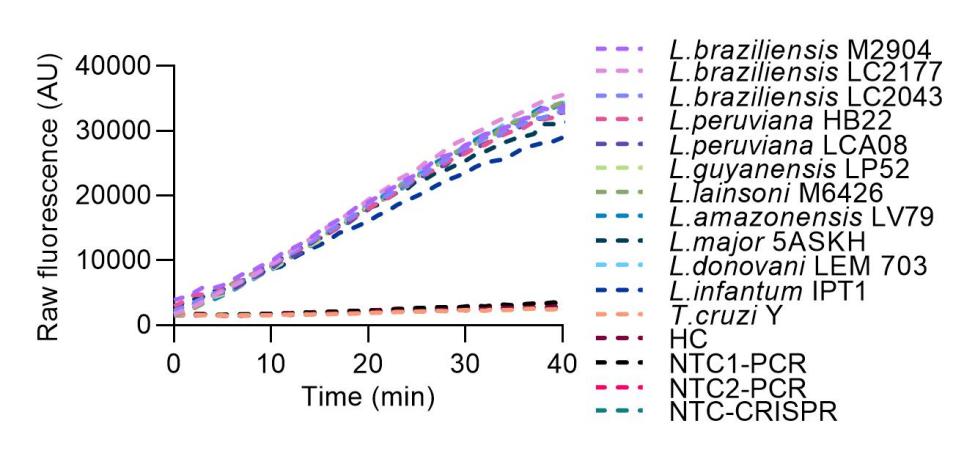

**F**

18S rDNA - primer set 3

**Figure S3.**

(**A-D**) Time course data of Cas12a detection experiments in Figure 4A. Two independent amplification (with kDNA primer set 1) and detection runs (**A**: first experiment, **C**: second experiment) were performed for the analytical specificity evaluation with DNA samples from 12 reference strains. (**A**, **C**) Raw fluorescence signal (arbitrary units, AU) over 2 h from Cas12a reactions. (**B**) Same dataset as in **A**, but zoomed to the first 40 min of the Cas12a assay. (**D**) Same dataset as in **C**, but zoomed to the first 40 min of the Cas12a assay. Based on the fluorescence kinetic curves, the data in this figure are used for detection analysis in Figure 4A at 20-min time point.

(**E-H**) Time course data of Cas12a detection experiments in Figure 4B. Two independent amplification (with 18S primer set 3) and detection runs (**E**: first experiment, **G**: second experiment) were performed for the analytical specificity evaluation with DNA samples from 12 reference strains. (**E**, **G**) Raw fluorescence signal (arbitrary units, AU) over 2 h from Cas12a reactions. (**F**) Same dataset as in **E**, but zoomed to the first 40 min of the Cas12a assay. (**H**) Same dataset as in **G**, but zoomed to the first 40 min of the Cas12a assay. Based on the fluorescence kinetic curves, the data in this figure are used for detection analysis in Figure 4B at 10-min time point.

Fluorescence measurements in this figure were made on the Synergy H1 plate reader.

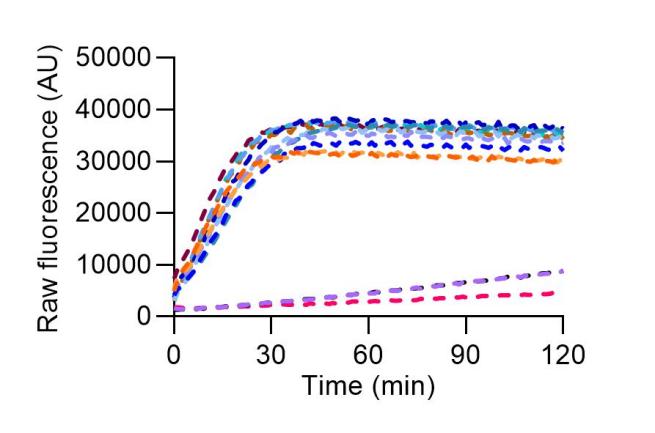

**E**

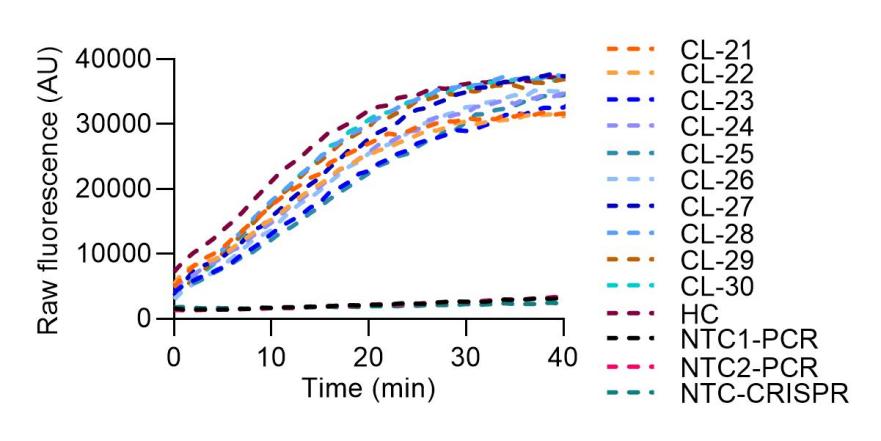

**F**

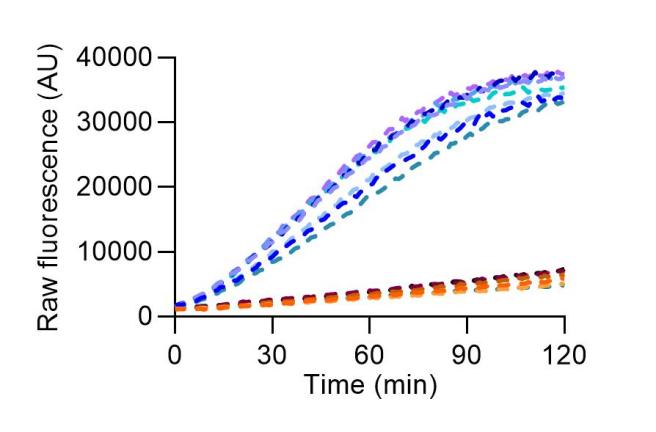

**A**

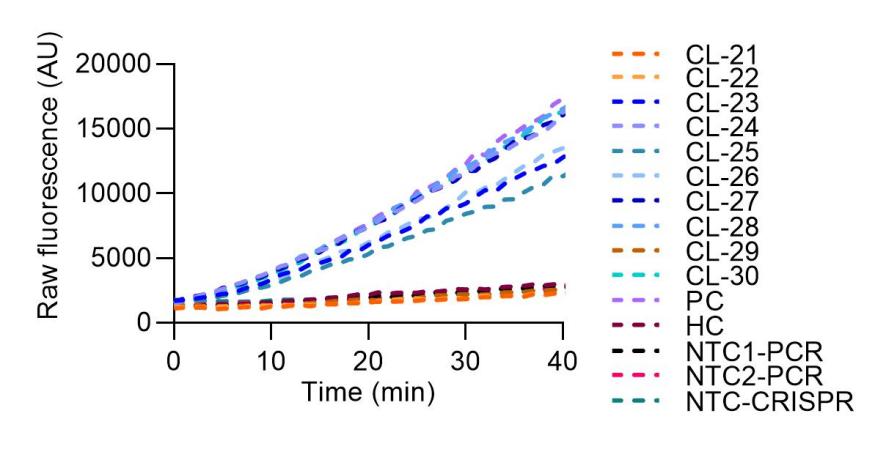

**B**

kDNA

18S rDNA

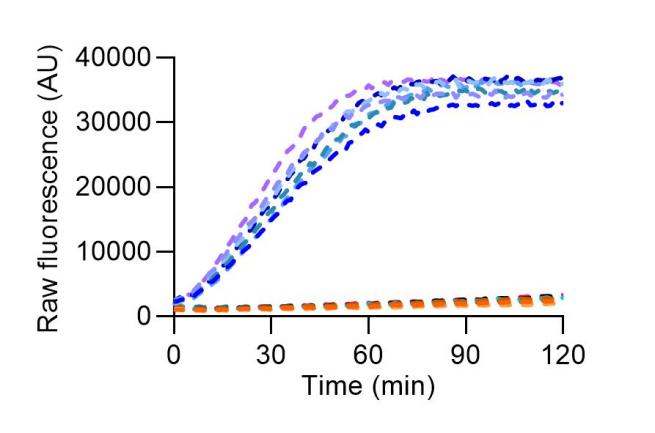

**C**

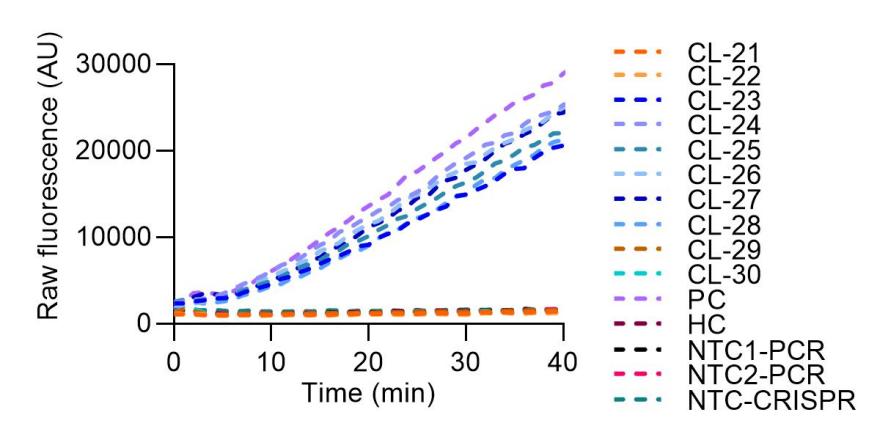

**D**

RNase P

**Figure S4.** Raw fluorescence curves generated by Cas12a detection of *Leishmania* kDNA (**A**) and 18S rDNA (**C**) targets, and a sample control, human RNase P gene (**E**), in clinical samples. A group of clinical samples (n = 10; coded CL-21 to CL-30) run in the same assay plate per target DNA is shown in this figure. Fluorescence measurements were made on the Synergy H1 plate reader. (**A**, **C**, **E**) Raw fluorescence signal (arbitrary units, AU) over 2 h from Cas12a reactions. (**B**) Same dataset as in **A**, but zoomed to the first 40 min of the Cas12a assay. (**D**) Same dataset as in **C**, but zoomed to the first 40 min of the Cas12a assay. (**F**) Same dataset as in **E**, but zoomed to the first 40 min of the Cas12a assay. Out of the 10 tested samples shown, 7 resulted in pronounced fluorescence curves indicating presence of *Leishmania* kDNA molecules (**A**), 6 showed robust fluorescence curves for the 18S rDNA target (**C**), and RNase P was detected in all of them (**E**).

kDNA

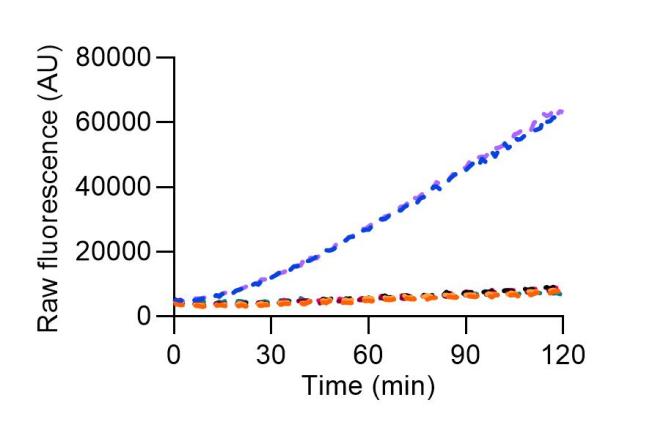

**A**

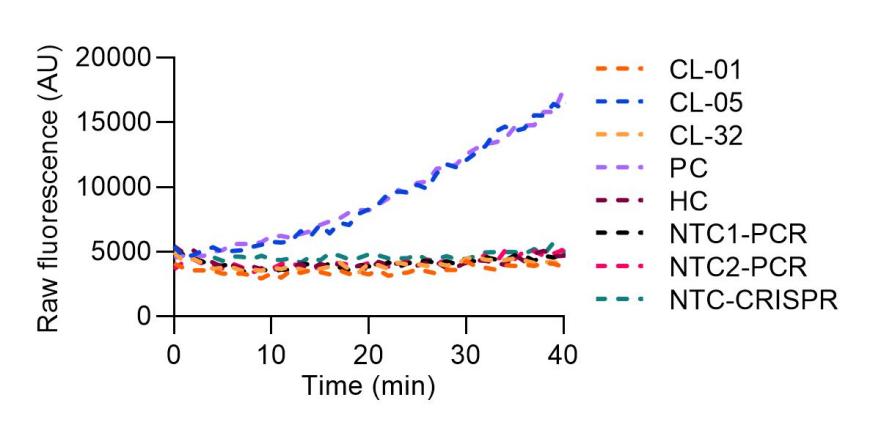

**B**

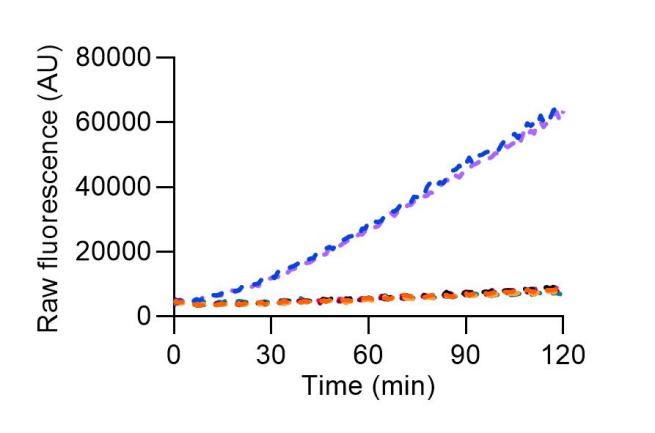

**C**

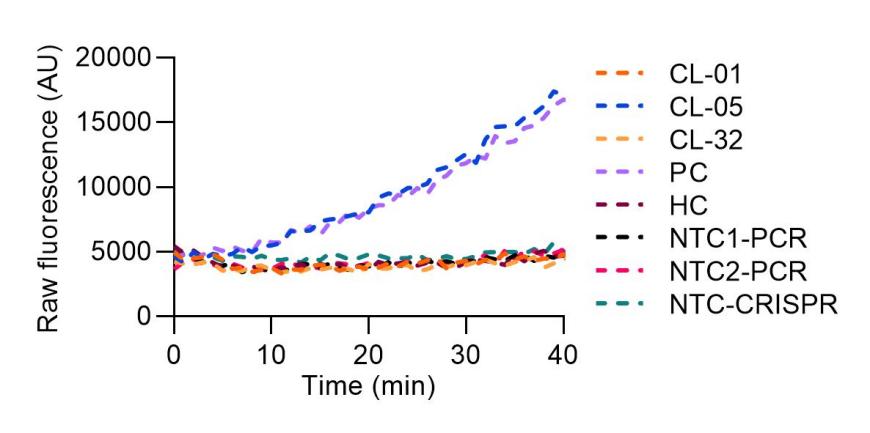

**D**

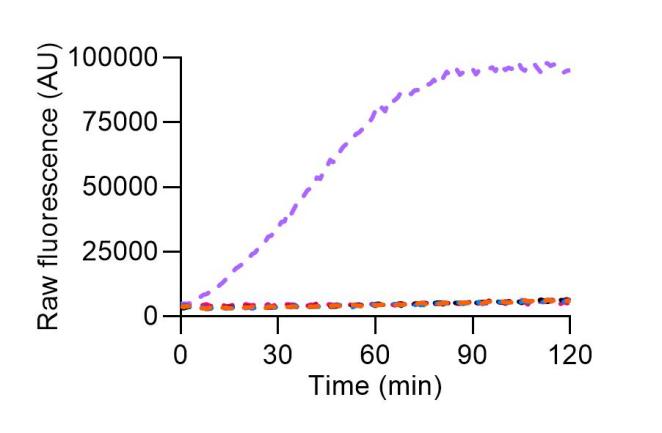

**E**

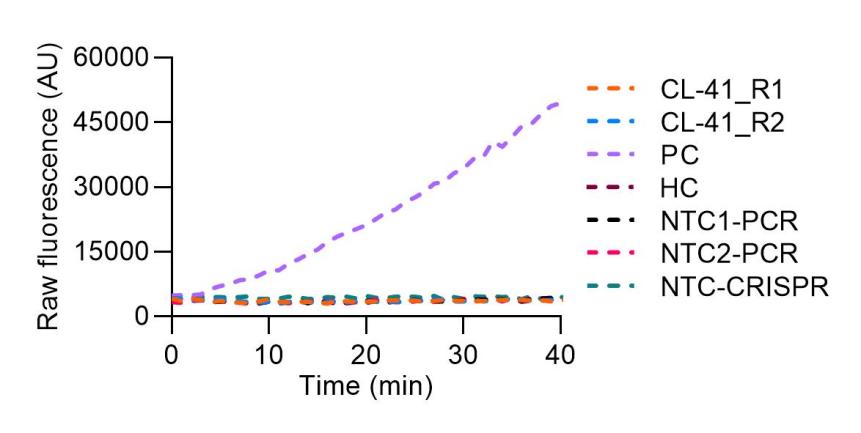

**F**

18S rDNA

**Figure S5.**

(**A, C**) Raw fluorescence curves generated by Cas12a detection of the *Leishmania* kDNA target in retested clinical samples, which had discordant results (samples CL-01 and CL-32) or low positive signal (sample CL-05) in the first measurements taken on the Synergy H1 plate reader. Duplicate PCR reactions were amplified in parallel with the kDNA primer set 1 (**A**: replicate 1, **C**: replicate 2) followed by Cas12a detection. Raw fluorescence signal (arbitrary units, AU) over 2 h from Cas12a reactions is shown. (**B**) Same dataset as in **A**, but zoomed to the first 40 min of the Cas12a assay. (**D**) Same dataset as in **C**, but zoomed to the first 40 min of the Cas12a assay.

(**E**) Raw fluorescence curves generated by Cas12a detection of the *Leishmania* 18S rDNA gene in the retested clinical sample CL-41, which had discordant results in the first measurements taken on the Synergy H1 plate reader. Duplicate PCR reactions were amplified in parallel with the 18S primer set 3 followed by Cas12a detection (both replicates are shown in this figure). (**F**) Same dataset as in **E**, but zoomed to the first 40 min of the Cas12a assay.

All fluorescence measurements in this figure were made on the Cytation 5 plate reader.

**Supplementary Material**

**Supplementary File S1**

Sequence alignment files. S_File1.1 contains the alignment of available *L.* (*V*.) *braziliensis* kDNA minicircle sequences. Files S_File1.2 and S_File1.3 contain the alignments of 18S rDNA and kDNA minicircle sequences, respectively, shown in Figure 2B. Alignments were performed using clustalo command from ClustalX package locally. Alignment files can be visualized using AliView software (https://ormbunkar.se/aliview/) or another alignment/genome viewer.

**Supplementary File S2**

This file contains the dataset generated during the current study that concerns qPCR and PCR/CRISPR data on tested clinical samples and the determination of threshold cutoff values for Cas12a-based assay interpretation. Fluorescence data from Cas12a assays presented here were taken on the Synergy H1 plate reader.

**Supplementary Table S1**

Primer and crRNA template sequences used in this study.
